# Suppression of nonsense mutations by small, cyclic peptides

**DOI:** 10.1101/2025.04.25.650553

**Authors:** Nanna Birkmose, Saleem Y. Bhat, Arpan Bhattacharya, Paul R. Hansen, Thomas G. Andersen, Sophia Hunter, Emilie T. Larsen, Yale E. Goldman, Barry S. Cooperman, Charlotte R. Knudsen

**Affiliations:** Department of Molecular Biology and Genetics, Aarhus University, DK-8000 Aarhus C, Denmark; Department of Chemistry, University of Pennsylvania, PA 19104, USA; Departments of Pharmacology and of Molecular and Cellular Biology, University of California, Davis, CA 95616, USA; Department of Drug Design and Pharmacology, University of Copenhagen, DK-2100 København Ø, Denmark; Interdisciplinary Nanoscience Center (iNANO), Aarhus University, DK-8000 Aarhus C, Denmark

## Abstract

Premature termination codons in mRNAs result from nonsense mutations and hinder the translation of full-length, functional proteins. Nonsense mutations cause numerous serious genetic diseases, including cystic fibrosis and Duchenne muscular dystrophy. Several small-molecule drugs have been reported that could potentially ameliorate these diseases by promoting translational readthrough at the premature termination codon. However, utilization of many of these molecules faces problems such as limited efficacy or high cellular toxicity. Using a selection strategy in *Saccharomyces cerevisiae* coupling suppression of endogenous nonsense mutations to cell survival, we identified ten readthrough-promoting cyclic peptides from a DNA-encoded library. The selected cyclic peptides suppress nonsense mutations in various reporter genes, and the candidates inducing the highest readthrough levels display no observable cytotoxicity in yeast. Mutational analysis of the most promising cyclic peptide demonstrate that most amino acid side chains contribute to the readthrough-stimulating activity. Importantly, this cyclic peptide appears to bind directly to the eukaryotic core translation machinery and promotes readthrough *in vitro* by interfering with ribosomal decoding. Our results suggest that small, cyclic peptides selected *in vivo* could represent a novel drug type to treat the many incurable human genetic diseases that are caused by nonsense mutations.

## INTRODUCTION

Nonsense mutations introduce premature termination codons (PTCs) in mRNAs and account for about 11% of all described mutations causing genetic disease in humans (1). Transcripts bearing PTCs are targeted for degradation via the nonsense-mediated mRNA decay (NMD) surveillance pathway (2,3). Yet, an estimated 5-30% of all PTC-containing transcripts are estimated to escape NMD (4), and thus produce truncated proteins due to premature termination of translation. Nonsense mutations typically lead to a very serious disease phenotype due to a complete loss of protein function (5). Examples include cystic fibrosis, Duchenne muscular dystrophy, and various cancers.

Translation termination is normally induced by the presence of a termination codon in the ribosomal A site. In eukaryotic cells, the release factor eRF1 recognizes the termination codon with high fidelity and triggers termination by peptidyl-tRNA hydrolysis (6,7), a process that is further stimulated by the GTPase eRF3 (8). Occasionally, a near-cognate tRNA binds the termination codon instead of eRF1 and inserts an amino acid in the growing polypeptide chain, thus causing the ribosome to continue translation (9,10). This process, termed translational readthrough, occurs naturally at an estimated low rate of 0.01-0.3% (11,12).

It was observed more than four decades ago that certain drugs promote the readthrough of PTCs in *S. cerevisiae* (13). Major efforts have since then gone into the development of readthrough-promoting compounds as a way of treating genetic diseases caused by nonsense mutations (14,15). Aminoglycoside antibiotics promote readthrough by binding the ribosome and promoting tRNA mispairing at PTCs (16,17) but are of limited therapeutic use due to frequently observed oto- and nephrotoxicity (18,19). A synthetic aminoglycoside derivative termed NB-124 or ELX-02 has been developed with higher efficacy and reduced cellular toxicity (20–22), but a recent phase II trial in cystic fibrosis patients reported no significant improvement upon treatment (15). Ataluren is a readthrough-stimulating oxadiazole molecule identified in a cell-based high-throughput screening (23). The compound promotes readthrough by inhibiting release factor activity (16). Ataluren has a good safety profile but showed limited and variable readthrough efficiency in phase III clinical trials (24,25). Nevertheless, ataluren has been conditionally approved for the treatment of Duchenne muscular dystrophy in the European Union and seems to work well for a subset of patients (26). Several other readthrough-promoting compounds have been reported over the years, but only ataluren has so far reached clinical approval (15). Clearly, new readthrough compounds are needed that target nonsense mutations with high efficacy and minimal cellular toxicity.

Small, cyclic peptides have potential as therapeutics. Their relatively large size makes them suitable for targeting large molecular surfaces such as protein-protein or protein-DNA/RNA interactions. Cyclization also makes the peptides more resistant to degradation and results in a more rigid conformation, allowing them to bind their target with high affinity and selectivity (27–29). Powerful methods have emerged to screen or select very large, combinatorial libraries of cyclic peptides for modulators of challenging targets (30–32). Here, DNA-encoded libraries are advantageous, since they can encode millions of variants with easy identification of potential hits by sequencing (33). Split-intein circular ligation of peptides and proteins, abbreviated SICLOPPS, allows for *in vivo* screening or selection for intracellular target binders (34). SICLOPPS libraries encode peptides flanked by intein segments (35) that facilitate a head-to-tail backbone cyclization after transcription and translation in cells (29). SICLOPPS has been widely used for early-stage drug discovery of cyclic peptides with therapeutic potential, such as a cyclic peptide inhibitor of hypoxia signaling with relevance in cancer (36) and cyclic peptides with antimicrobial activity (37). We recently reported the construction and characterization of a *S. cerevisiae* SICLOPPS library encoding randomized cyclic hexapeptides (38). The library was shown to be correctly encoded, and only marginal changes in library composition were observed upon expression in yeast, paving the way for unbiased selection studies of cyclic peptides (38).

Here, we report a novel SICLOPPS-based selection system *in S. cerevisiae*, coupling cell survival to the expression of cyclic peptides capable of promoting readthrough of reporter genes carrying nonsense mutations. Using this approach, we identified ten cyclic peptides capable of suppressing nonsense mutations in various genetic contexts. Notably, four of the most promising candidates (named CP06, CP07, CP35, and CP55) displayed no observable toxicity in yeast cells. Alanine-scanning mutagenesis of CP55 revealed that most amino acids in the cyclic peptide are part of the functional motif. Moreover, mutational analysis also revealed that cyclization is crucial for the readthrough activity of CP55. *In vitro* studies of CP55 in a eukaryotic translation system revealed a mechanism of action targeting the core translation machinery. Together, our study highlights the possibility of using cyclic peptides in the development of therapies to treat diseases caused by nonsense mutations.

## MATERIALS AND METHODS

### SICLOPPS library construction

A yeast SICLOPPS library was prepared as previously described (38). In brief, intein gene segments from *Synechocystis* sp. PCC6803 (35) were cloned into the pESC-LEU yeast epitope tagging vector (Agilent Technologies #217452) in a reversed order, with positioning to add a c-myc tag to the C-terminus of intein N. The randomized sequence encoding the cyclic peptides (TGC-(NNS)_5_; N = A, T, G, C and S = G, C) was introduced between the intein segments in PCR reactions using a degenerate library primer. The resulting PCR fragments were used as DNA templates in subsequent “zipper” PCR reactions to remove potential mismatches (39). Purified PCR products and the intein-containing vector were digested with restriction enzymes, ligated, and purified by drop dialysis (39). The ligated products were transformed by electroporation into highly competent *Escherichia coli* SIG10 cells (Sigma-Aldrich) (38) and plated on large dishes of LB agar added 100 µg/mL ampicillin. The SICLOPPS library was obtained by isolation of plasmid DNA from bacterial colonies collected by scraping. The resulting library was estimated to hold 8 × 10^7^ individual library members (38). The empty vector, used throughout the study as a negative control, refers to pESC-LEU with insertion of the intein gene segments.

### Large-scale library transformation in *S. cerevisiae*

*S. cerevisiae* IS110-18A (([PSI]^−^ *MATa aro7-1 leu2-2 ilv1-2 his4-166 lys2-101 met8-1 trp5-48 ura4-1*) (40) cells were transformed with the constructed SICLOPPS library using a large-scale method employing lithium acetate and heat shock (41). An overnight stationary-phase culture in yeast peptone dextrose (YPD) medium was used to prepare a 330-mL culture in YPD starting at an OD_600_ of 0.2 and incubated at 30°C with shaking until reaching exponential-phase growth (OD_600_ ∼0.6-0.7). Cells were harvested by centrifugation, washed in Tris-EDTA (TE) buffer (10 mM Tris-HCl, 1 mM EDTA, pH 7.5), and resuspended in 1.5 mL TE containing 0.1 M lithium acetate. 1 mL of cells were added to a tube containing 20 µg SICLOPPS library plasmid DNA and 2 mg boiled salmon testes single-stranded carrier DNA (Sigma-Aldrich). The DNA was mixed with 6 mL TE and 0.1 M lithium acetate in 40% PEG 4000, thoroughly mixed by vortexing, and incubated for 30 minutes at 30°C with shaking. Then, 700 µL dimethyl sulfoxide (DMSO) was added to the cells and gently mixed by turning the tube. The cells were heat-shocked for 15 minutes at 42°C in a water bath and allowed to recover in 10 mL YPD for one hour at 30°C. Subsequently, the cells were harvested, washed in TE buffer, resuspended in TE buffer, and plated on a large (245 × 245 mm) dish of synthetic defined (SD)/–Leu agar added 2% glucose. A small volume of cells was retained, serially diluted, and plated separately to estimate the number of transformants. The plates were wrapped in parafilm and left for up to three days in a 30°C incubator.

### Selection in *S. cerevisiae* and isolation of candidate library members

*S. cerevisiae* colonies transformed with the cyclic peptide library were collected by scraping and harvested in TE buffer by centrifugation. The cell suspension was washed in TE buffer and resuspended in TE buffer. Cell counts were estimated via OD_600_ measurements. Following this, 800,000 cells were plated on 24 140-mm Petri dishes of SD/–Leu/–Met agar added 2% galactose. The same number of cells were also plated on the following control plates: SD/–Leu agar + 2% galactose, SD/–Leu agar + 2% glucose, SD/–Leu/–Met agar + 2% glucose, and SD/–Leu/–Met agar + 2% glucose + 200 µg/mL G418 (Sigma-Aldrich). The plates were wrapped in parafilm and placed in a 30°C incubator. Cell growth was monitored daily for five days. Representative images were taken on separate days to enable a comparison of similar growth stages of cells (Table S1). Yeast colonies growing on the selective medium were picked and individually cultured in SD/–Leu added 2% glucose at 30°C with shaking until reaching the stationary phase, and total DNA was extracted for each culture using a yeast DNA extraction kit (Thermo Scientific). Plasmid DNA was amplified by electroporation of total yeast DNA into *E. coli* SIG10 cells. Plasmid DNA was extracted from separate overnight cultures of transformed *E. coli* cells grown in LB + 100 µg/mL ampicillin at 37°C with shaking, using the GeneJET Plasmid Miniprep Kit (Thermo Scientific).

### Spot assays

Purified plasmid constructs expressing candidate readthrough-promoting cyclic peptides or mutants thereof were individually transformed into *S. cerevisiae* IS110-18A cells using standard lithium acetate treatment and heat shock. Colonies of transformed cells were cultured overnight in SD/–Leu added 2% glucose at 30°C with shaking. Cultures were diluted in sterile H_2_O or the appropriate selective medium to an OD_600_ of 1.0. Dilutions with adjusted cell counts were then further serially ten-fold diluted in sterile H_2_O and spotted in 5-µL droplets on selective medium and control agar plates. The plates were wrapped in parafilm and incubated at 30°C with growth being monitored daily. Representative images were taken on separate days to enable a comparison of similar growth stages of cells (Table S1). The ability of cyclic peptides to suppress nonsense mutations was evaluated by comparing cells grown on the selective medium expressing cyclic peptides to cells expressing an empty vector or expressing a random cyclic peptide (i.e. randomly picked and with no phenotypic effect; c(Cys-Ser-Gly-Val-Leu-Glu) (38)). The sequences of cyclic peptide constructs enhancing cell growth were determined by Sanger sequencing (Eurofins Genomics TubeSeq) (Sequencing primers are listed in Table S2).

### Western blotting

*S. cerevisiae* IS110-18A cells were transformed with plasmid constructs expressing readthrough-promoting cyclic peptides or mutants thereof using standard lithium acetate treatment and heat shock. Controls expressing an empty vector or random cyclic peptides (i.e. randomly picked and with no phenotypic effect; (Cys-Ser-Gly-Val-Leu-Glu) and c(Cys-Leu-Trp-Thr-Leu-Gly) (38)) were included. Colonies of transformed cells were cultured until reaching the stationary phase in SD/–Leu added 2% galactose at 30°C with shaking. Cultures were chilled on ice, pelleted by centrifugation, and washed in ice-cold H_2_O. Cell pellets were frozen in liquid nitrogen and stored at −70°C. Cell lysis was performed by resuspension in a lysis buffer (0.1% SDS, 1% NP-40, 0.5% sodium deoxycholate, 50 mM Tris-HCl pH 8.0, 150 mM NaCl, 0.57 mM (100 µg/mL) phenylmethylsulfonyl fluoride, 1× cOmplete EDTA-free protease inhibitor cocktail (Roche)) followed by rapid freeze-thawing, using 1 mL of lysis buffer per 10^9^ cells (42). Aliquots of lysed cells were mixed with an SDS loading buffer. Proteins were separated by SDS-PAGE, transferred to a PVDF membrane, and the membrane was analyzed by Western blotting applying standard procedures. The antibodies used were a primary antibody, mouse anti-c-myc (Sigma-Aldrich #M4439) diluted 1:1000, and a secondary antibody, horse radish peroxidase (HRP)-conjugated goat α-mouse (Dako) diluted 1:5000.

### Growth assay in liquid culture

Colonies of *S. cerevisiae* IS110-18A cells transformed with readthrough-promoting cyclic peptides were pre-incubated in 5 mL SD/–Leu added 2% galactose at 30°C with shaking for 24 hours to ensure that cells were already expressing cyclic peptides at the start of the growth assays. The cells were washed in sterile H_2_O and used to set up 15-mL cultures in non-selective SD/–Leu + 2% galactose or readthrough-selective SD/–Leu/–Met + 2% galactose medium with a starting OD_600_ of 0.15. Growth of the cultures was monitored over several days by measuring OD_600_ on suitable cell culture dilutions and comparing to the growth of cells expressing a random cyclic peptide (i.e. randomly picked and with no phenotypic effect; c(Cys-Ser-Gly-Val-Leu-Glu) (38)).

### Site-directed mutagenesis and alanine scanning

The SICLOPPS library-derived plasmid expressing CP55 (c(Cys-Tyr-Phe-Ser-Val-Gly)), specifically pESC-LEU with insertion of the CP55-encoding sequence flanked by intein domains promoting cyclization, was used as a DNA template for mutagenesis. Mutations introduced individually in CP55 were C1S, Y2A, Y2W, F3A, V4A, S5A, and G6A using the Quikchange Lightning Site-Directed Mutagenesis kit (Agilent Technologies) and mutagenic primers (Table S2). Mutations in intein N (T69A+H72A), blocking the first step in intein processing (43), or mutations in intein C (H24L+F26A), blocking the final processing step leading to a release of cyclized peptide (44), were introduced using a modified Quikchange protocol with primers containing extended non-overlapping segments (45) (Table S2). Vectors with confirmed mutations were re-transformed into *E. coli*, followed by repeated plasmid purification and Sanger sequencing for double verification of the relevant mutations.

### Chemical synthesis of cyclic peptide CP55

The readthrough-promoting cyclic peptide CP55 (c(Cys-Tyr-Phe-Ser-Val-Gly)) was synthesized by 9-fluorenylmethyloxycarbonyl (Fmoc) solid-phase peptide synthesis on a 2-chloro-trityl resin (0.1 mmol) preloaded with the first amino acid. Chain elongation was achieved with single couplings using 3.5 equivalents each of Fmoc-*N^α^*-L-amino acid, 1-hydroxy-7-azabenzotriazole (HOAt), and hexafluorophosphate azabenzotriazole tetramethyl uranium (HATU) and 7 equivalents of diisopropylethylamine (DIEA). After 2 hours, the resin was washed with dimethylformamide (DMF) (3 × 1 min), dichloromethane (DCM) (3 × 1 min), and DMF again (5 × 1 min). The Fmoc group was removed by treatment with piperidine–DMF (1:4) (3 × 1 min); followed by washing as described above. The protected peptide was cleaved from the resin with hexafluoroisopropanol–DCM (1:4), (3 × 30 min), the filtrate was collected and evaporated to dryness. In-solution cyclization C->N terminus was carried out using 7-azabenzotriazol-1-yloxy)tripyrrolidinophosphoniumhexafluoro-phosphate (PyAOP) (1 equivalent) in the presence of HOAt (1 equivalent)/DIEA (2 equivalents) in DMF–DCM (6:94) (46). Following overnight reaction, DCM was evaporated, and water was added to precipitate the peptide. The peptide was then lyophilized and treated with trifluoroacetic acid (TFA):triisopropylsilane:H_2_O (95:2.5:2.5) for 2 hours. TFA was evaporated, the peptide precipitated in ether, lyophilized, and characterized. The main product was the desired cyclic peptide c(Cys-Tyr-Phe-Ser-Val-Gly) (calculated: 657.76 found: 657.68). The cyclic peptide was dissolved in pure Milli-Q water at a concentration of 2.5 or 5 mM, aliquoted, quick-frozen in liquid nitrogen, and stored at −80°C.

### Components for *in vitro* translation assays

Eukaryotic 80S ribosomes from brine shrimp (*Artemia Salina*) cysts and carrier *E. coli* 70S ribosomes were isolated and purified as previously described (16,47). Yeast translation elongation factors eEF1A and eEF2 and human release factors eRF1 and eRF3 were purified as reported earlier (16,48). Yeast Phe, Arg, and Trp tRNAs along with *E. coli* Lys, Val, and Gln tRNAs were isolated from bulk yeast and *E. coli* tRNAs, respectively, by hybridization to complementary, biotinylated DNA oligos (49). The tRNAs were charged with their cognate amino acids in an aminoacylation assay using crude yeast or *E. coli* tRNA synthetases for yeast and *E. coli* tRNAs, respectively (50,51). RNA transcripts, containing the reporter sequences and a cricket paralysis virus internal ribosome entry site (IRES), were prepared by *in vitro* transcription of plasmid variants produced commercially (Twist Biosciences).

To prepare POST5 complexes, *A. salina* 80S ribosomes were incubated with either Trp-IRES (encoding Phe-Lys-Val-Arg-Gln-Trp) or Stop-IRES (encoding Phe-Lys-Val-Arg-Gln-Stop) RNAs at 37°C for 30 minutes in Buffer 4 (40 mM Tris-HCl (pH 7.5), 80 mM NH_4_Cl, 5 mM magnesium acetate, 100 mM potassium acetate, 3 mM β-mercaptoethanol). The resulting 80S-IRES complexes were ultracentrifuged through a 1.1 M sucrose solution in Buffer 4 (1.6× of reaction volume) at 540,000 × g for 90 min at 4°C, and the pellet was dissolved in Buffer 4. Following this, Trp-POST5 and Stop-POST5 complexes were made by incubating 80S-IRES complexes with charged tRNAs, elongation factors eEF1A and eEF2, and 1 mM GTP as previously described (52), thus producing 80S ribosomes with Phe-Lys-Val-Arg-Gln-tRNA^Gln^ in the P site and a UGG (Trp-POST5) or UGA (Stop-POST5) codon in the A site. POST5 concentration was estimated from both A_260_ measurements and the stoichiometry of radioactively labeled pentapeptide per 80S ribosome as previously reported (52).

### Readthrough assay based on co-sedimentation

Translational readthrough was quantified *in vitro* by co-sedimentation of radioactively labeled hexapeptides with ribosomes, formed by the insertion of a [^3^H]-labeled near-cognate Trp-tRNA^Trp^ at a UGA stop codon in the Stop-POST5 complexes (16,53). Two solutions were prepared. One solution contained the POST5 complex in Buffer 4 with GTP added, while the other solution contained [^3^H]-Trp-tRNA^Trp^, elongation factors eEF1A and eEF2, release factors eRF1 and eRF3, and GTP in Buffer 4. CP55 was added to both solutions. The reactions were initiated by thoroughly mixing the two solutions at room temperature. After 90 seconds, the reactions were quenched by adding an excess of 0.5 M ice-cold MES buffer (pH 6.0) and placing the reactions on ice. The final concentrations of components in the reactions were 0.05 µM POST5 complex, 1 mM GTP, 0.2 µM [^3^H]-Trp-tRNA^Trp^, 1 µM eEF1A, 1 µM eEF2, 0.04 µM eRF1, 0.1 µM eRF3, and varying concentrations of CP55. Where indicated, eRF1 and eRF3 were omitted. Ribosomes were pelleted by first adding 100 pmol *E. coli* 70S carrier ribosomes, followed by ultracentrifuging through a 1.1 M sucrose solution in Buffer 4 at 540,000 × g for 70 min at 4°C. The supernatant was decanted, and the ribosomal pellet quickly washed in Buffer 4. Following resuspension in Buffer 4, [^3^H] radioactivity was measured to determine tryptophan incorporation at the UGA stop codon. As a positive control, tryptophan insertion at its cognate UGG codon in Trp-POST5 complexes was likewise measured.

### Termination assay based on fluorescence anisotropy

Translation termination was quantified by the release of a fluorescently labeled hexapeptide from the POST5 complex, resulting in changes in fluorescence anisotropy (52). The pentapeptide, coupled to the P-site tRNA in the POST5 complexes, was labeled by utilizing Atto 647-labeled Lys-tRNA^Lys^ during POST5 complex formation (52). Two solutions were prepared. One solution contained the POST5 complex in Buffer 4 with GTP added, while the other solution contained release factors eRF1 and eRF3 in Buffer 4 containing GTP. CP55 was added to both solutions. The final concentrations of components in the reactions were 0.05 µM POST5 complex, 1 mM GTP, 0.13 µM eRF1, 1.6 µM eRF3, and varying concentrations of CP55. The reactions were initiated by simultaneously mixing the two solutions for all reactions using a multichannel pipette in a 384-well flat-bottom plate (CORNING). Real-time monitoring of the release of Atto 647-labeled peptide from the POST5 complexes through changes in anisotropy was conducted at 25°C using a TECAN SPARK multimode reader equipped with a monochromator. Fluorescence anisotropy decay traces were processed with GraphPad Prism, using a one-phase exponential decay model to obtain rates of peptide release.

### Readthrough assay by single-molecule FRET

Translational readthrough was quantified in an assay monitoring the efficiency of fluorescence resonance energy transfer (FRET) between the ribosomal P-site tRNA and a near-cognate tRNA bound to a UGA stop codon in the A site by single-molecule total internal reflection fluorescence (TIRF) microscopy (53,54). All single-molecule TIRF studies were conducted at 24°C. An enzymatic oxygen scavenging system to reduce photobleaching consisted of 2 mM protocatechuic acid (PCA), 50 nM protocatechunate 3,4 dioxygenase (PCD), 1 mM cyclooctatetraene (COT), 1 mM 4-nitrobenzyl alcohol (NBA), and 1.5 mM 6-hydroxy-2,5,7,8-tetramethyl-chromane-2-carboxylic acid (Trolox) (all from Sigma-Aldrich) and was used for all dilutions, complex formation, and single-molecule imaging. Image stacks were recorded using a custom-built, objective-type TIRF microscope, based on a commercial inverted microscope (Eclipse Ti-E, Nikon), at a frame rate of 10 s^−1^ (54). The microscope could perform alternating-laser excitation (ALEX) between 532 nm and 640 nm laser beams using an acousto-optic tunable filter (AOTF) to switch wavelengths.

Biotinylated Stop-POST5 or Trp-POST5 complexes, containing Phe-Lys-Val-Arg-Gln-tRNA^Gln^(Cy5) in the P site, and a UGA stop codon or a UGG codon in the A site for Stop-POST5 and Trp-POST5, respectively, were injected into the streptavidin-coated, PEGylated slide chamber. After a 5-minute incubation, excess unbound ribosomes were washed out. Then, a ternary complex consisting of Trp-tRNA^Trp^(Cy3), eEF1A, and GTP was injected in the presence or absence of CP55 and incubated for 5 mins to form the Stop-PRE6 or Trp-PRE6 complex, and subsequently, excess reagents were flowed out and movies were recorded. Trp-PRE6 or Stop-PRE6 formation, occurring when Trp-tRNA^Trp^(Cy3) was accommodated into the A site, was confirmed by a tRNA-tRNA FRET signal (FRET efficiency ≈ 0.5) between tRNA^Gln^(Cy5) in the P site and the peptidyl-tRNA, Phe-Lys-Val-Arg-Gln-Trp-tRNA^Trp^(Cy3), in the A site. The recorded movies were analyzed by a custom-made software program developed as an ImageJ plugin (http://rsb.info.nih.gov/ij), and in Python. Error bars represent the mean ± SEM, which is calculated from the total number of spots.

## RESULTS

### Identification of ten novel, nonsense mutation-suppressing cyclic peptides

In this study, our recently reported yeast SICLOPPS library (38) was applied for selection in *S. cerevisiae*. The DNA-encoded cyclic peptide library was constructed in the pESC-LEU vector, allowing for inducible expression of peptides by galactose and the use of a *LEU2* marker to select transformed cells in the absence of leucine. The library encodes a fusion of random peptides with flanking, cyclizing intein fragments (34,39), and a c-myc tag located in the C-terminus of the fusion protein. The cyclic peptides are encoded by five randomized codons, each specifying one of the 20 canonical amino acids, and a cysteine, which facilitates cyclization (34). The SICLOPPS library consists of 8 × 10^7^ library members and encodes the 3.2 million possible amino acid combinations. Moreover, the library was shown to be expressed and cyclized in yeast (38).

Potential cyclic peptides with the capacity to promote translational readthrough of nonsense mutations were identified in a large-scale selection from the cyclic peptide library after transformation into the *S. cerevisiae* strain IS110-18A (40). This strain contains nonsense mutations in different essential biosynthesis genes and has previously been utilized to discover small molecules that potentiate the readthrough effect of aminoglycoside antibiotics (40). These nonsense mutations hinder the expression of full-length proteins due to premature translation termination at the resulting PTCs, causing the essential biosynthesis proteins to be non-functional. Thus, the cell is unable to grow in a selective medium lacking the relevant metabolite (Figure 1A). Nonsense-mutation suppression, as induced by a readthrough-inducing cyclic peptide promoting near-cognate tRNA binding in the ribosomal A site, restores protein function and thereby promotes cell survival (Figure 1A). Specifically, the selection was conducted against the *met8-1* nonsense mutation, which converts the glutamate codon GAG to a UAG termination codon and is followed by an adenine (40). The UAG-A sequence was chosen, since it theoretically provides mid-range ease of promoting readthrough at a given stop codon context (11,55).

**Figure 1.**
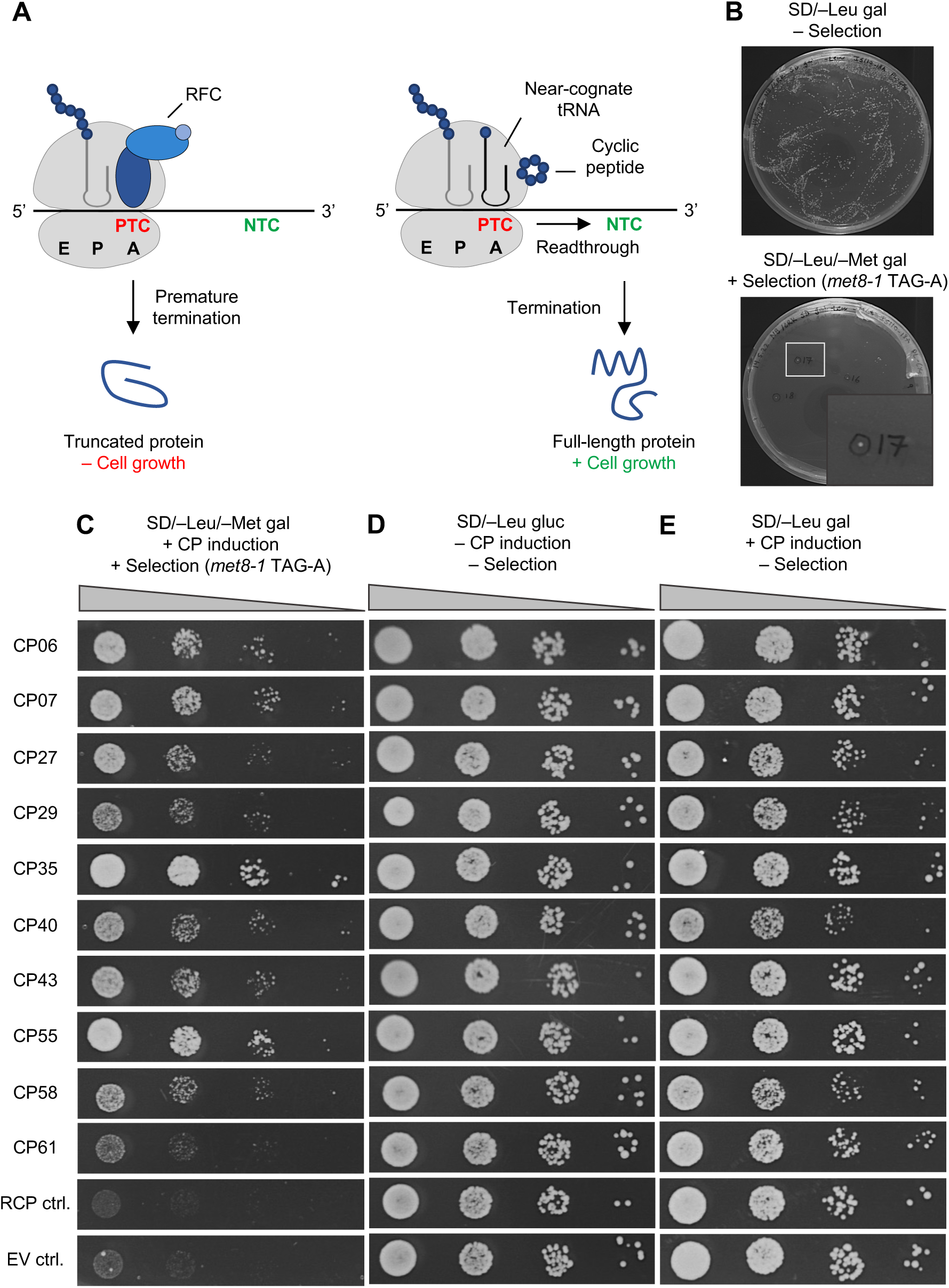
Selection in *S. cerevisiae* results in ten cyclic peptides suppressing nonsense mutations. (**A**) Outline of the selection principle. A genomic reporter gene contains a premature termination codon (PTC). The reporter gene is essential for cell growth on the selective medium. Upon translation, truncated and non-functional proteins are produced due to premature translation termination mediated by the release factor complex (RFC), and the cells are unable to grow (left part). Expression of a cyclic peptide (CP) that promotes translational readthrough will result in amino acid insertion by a near-cognate tRNA, thus allowing translation to continue to the natural termination codon (NTC), restoring full-length protein and promoting cell survival on the selective medium (right part). (**B**) Representative images of hit colony identification. *S. cerevisiae* IS110-18A cells transformed with the SICLOPPS library were grown on plates containing selective medium (lower; representative example) or on a control plate (upper). The presence of galactose (gal) in the medium induces CP expression. The omission of leucine (–Leu) selects for transformed cells, while the omission of methionine (–Met) selects for cells expressing CPs capable of suppressing a nonsense mutation present in the *met8* (*met8-1* TAG-A) gene. Images were taken after five days (lower) or three days (upper) of incubation. (**C-E**) Validation of ten readthrough-promoting CPs after isolation and re-transformation of the encoding constructs into fresh *S. cerevisiae* cells. The resulting transformants were spotted in serial ten-fold dilutions. (**C**) Growth enhancement on selective medium through nonsense-mutation suppression was evaluated by comparison to cells expressing a random cyclic peptide (RCP) or an empty vector (EV). (**D**) Growth on non-selective, glucose-containing medium (no expression of CPs). (**E**) Growth on non-selective, galactose-containing medium (CP expression induced), with decreased growth indicative of cytotoxicity mediated by the expressed CP. Images were taken after four days (C), two days (D), or three days (E) of incubation. All spot assay images (C-E) are representative of five independent experiments.

The selection was conducted by large-scale transformation of *S. cerevisiae* IS110-18A cells with the SICLOPPS library, yielding 4.3 million independent transformants each encoding a cyclic peptide variant. Colonies of transformed cells were collected and re-plated on several agar plates of selective medium, to ensure the representation of all transformants, along with relevant controls. The selection medium contained galactose as a carbon source to induce cyclic peptide expression and lacked leucine and methionine to select for transformed cells and suppressors of the *met8-1* nonsense mutation, respectively. A total of 63 surviving colonies were identified on selective medium (a representative plate is shown in Figure 1B). Limited colony formation was observed on a glucose-containing control selective plate (Figure S1A), with colonies appearing only after a prolonged period of incubation. This indicates the necessity of library induction for the identification of surviving cells. Moreover, the addition of the readthrough-inducing aminoglycoside G418 (56) to the medium could independently induce colony formation, demonstrating that cell survival can be mediated by promoting readthrough on the selective medium (Figure S1B).

The library plasmids expressed in individual, surviving colonies were obtained by individually culturing the cells, extracting DNA, and amplification of plasmid DNA in *E*. *coli* after transformation with the extracted, total yeast DNA. For ten colonies, plasmid DNA extraction failed. The remaining 53 plasmid candidate library members were individually re-transformed into fresh IS110-18A cells and spotted on a selective medium lacking methionine to validate the growth-promoting effect of the expressed, cyclic peptides (Figure S2). Enhanced cell growth by PTC readthrough was assessed by comparison to cells expressing a random cyclic peptide (RCP) or an empty vector (EV), both of which should be unable to grow on a selective medium. The validation resulted in the identification of ten library members reproducibly suppressing nonsense mutations to varying degrees (Figure 1C). The remaining colonies were likely false positives that arose from spontaneous genomic mutations causing survival unrelated to cyclic peptide expression, a phenomenon that is not uncommon in yeast selections (41). The strongest nonsense suppressors, as judged by enhanced cell growth, were CP06, CP07, CP35, and CP55 (Figure 1C). A comparison of cells grown on non-selective medium in the presence of either glucose (no expression of cyclic peptides; Figure 1D) or galactose (expression of cyclic peptides; Figure 1E) could reveal any effects of the readthrough-stimulating cyclic peptides on general cell growth. Interestingly, this comparison revealed that some of the identified readthrough-promoting cyclic peptides (CP27, CP29, CP40, CP58, and CP61) caused general growth inhibition, while the expression of others did not affect cell growth (Figure 1E). Overall, ten candidate readthrough-promoting cyclic peptides of varying potency were discovered in the yeast selection, of which some displayed no observable cytotoxicity.

### Readthrough-promoting cyclic peptides can suppress several endogenous nonsense mutations

As the identified readthrough-stimulating cyclic peptides are derived from a SICLOPPS library, the peptides are expressed from DNA constructs transformed into cells followed by intein-mediated cyclization (34). Western blotting was applied to verify intein processing, thereby indirectly indicating the presence of cyclized peptides in the cells (57). Processing to release the peptide in a cyclized form results in the cleavage of intein C (4 kDa) and intein N (15.5 kDa), which are encoded in a reversed order in SICLOPPS libraries to promote cyclization (34). Thus, processing can be monitored by means of a c-myc-tag, located in the C-terminus of the fusion construct (Figure 2A) (39). *S. cerevisiae* strain IS110-18A cells were individually transformed with each of the ten readthrough-promoting cyclic peptide constructs, and lysates of cells grown to stationary phase were analyzed by Western blotting to detect expression and processing. All unprocessed intein-peptides could be detected (Figure 2B, upper band; unprocessed blot Figure S3A). Expression levels were generally high, albeit the growth-inhibiting cyclic peptides (Figure 1E) displayed reduced expression levels. Processing to release cyclized peptides could be detected for all candidates except CP58, which was generally expressed at a lower level (Figure 2B, lower band). Of note, high levels of processing did not seem to be the only determinant of readthrough potential, as CP55 showed very strong nonsense suppression but relatively weak processing.

**Figure 2.**
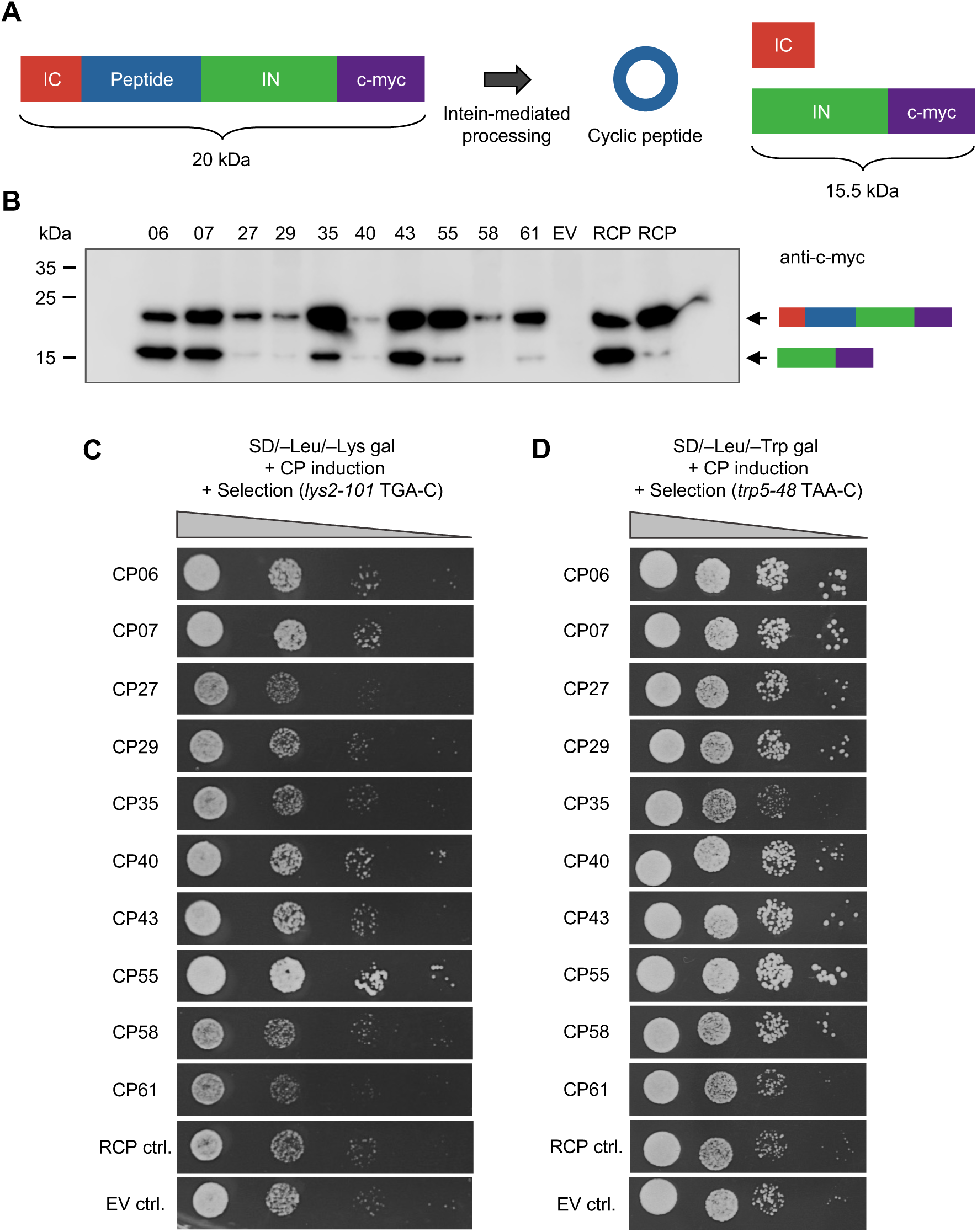
Readthrough-promoting cyclic peptides are produced in yeast and can suppress endogenous nonsense mutations. (**A**) Schematic depicting intein-mediated processing to produce cyclic peptides (CPs) intracellularly. Processing can be followed by Western blotting detecting a reduction in size, from 20 to 15.5 kDa, of the c-myc tagged intein N (IN), due to the release of intein C (IC) and the cyclized peptide from IN. (**B**) Western blotting detects the expression and processing of the intein-peptide fusion protein by means of an anti-c-myc antibody. Lysates of *S. cerevisiae* IS110-18A cells transformed with constructs encoding the readthrough-stimulating CPs were used, together with cells expressing an empty vector (EV), encoding intein segments without insertion of randomized sequences resulting in a frameshift, or two different random cyclic peptides (RCPs) as controls. The identities of the bands are shown as cartoons using the color code in panel A. (**C, D**) Constructs encoding the readthrough-promoting CPs were isolated, re-transformed into fresh *S. cerevisiae* IS110-18A cells, and spotted in serial ten-fold dilutions on a selective medium. The presence of galactose (gal) induces CP expression, while the omission of leucine (–Leu) selects for transformed cells. The additional omission of lysine (– Lys) (**C**) or tryptophan (–Trp) (**D**) selects for cells expressing CPs capable of suppressing nonsense mutations in the *lys2* (*lys2-101* TGA-C) and *trp5* (*trp5-48* TAA-C) genes, respectively. Growth enhancement through nonsense mutation suppression was evaluated by comparison to cells expressing RCP or EV as negative controls. Images were taken after four days of incubation and are representative of three independent experiments.

The ten identified readthrough-promoting cyclic peptides were further investigated for their ability to suppress PTCs in other reporter genes in the IS110-18A yeast strain (40). To this end, cultured cells expressing the candidate peptides were spotted on a selective medium without the relevant, essential metabolite produced by the protein encoded by the PTC-affected reporter gene. Several of the cyclic peptides could suppress TGA and TAA nonsense mutations, as evident by suppression in the *lys2-101* (TGA-C) and *trp5-48* (TAA-C) genes (Figures 2C and D). However, a high level of non-specific, background growth of cells on the selective medium (see RCP and EV controls in Figures 2C and D) likely masked any readthrough effect of the weakly acting cyclic peptides. The ten cyclic peptides were also tested for nonsense mutation suppression in the *ura4-1* (TAA-A) gene, but no growth was observed for any of the peptides (Figure S4A). The addition of the readthrough-stimulating compound G418 did likewise not result in any cell growth (Figure S4B), underlining the potential difficulty in inducing readthrough at this particular sequence (58,59). Moreover, attempts to perform a SICLOPPS library selection for suppressors of the *ura4-1* nonsense mutation failed to identify any candidates (data not shown). A nonsense mutation in the *ilv1-2* (TAA-C) reporter gene could only be suppressed by CP55 (Figure S4C). Of note, CP35 showed a very strong nonsense-mutation suppression in *met8-1*, but no observable activity for any other reporter genes tested (Figures 1C, 2C, 2D, S4A, and S4C).

Sequences of the ten readthrough-promoting cyclic peptides are shown in Table 1, as determined by Sanger sequencing of their encoding constructs. Listed is also a qualitative assessment of their cytotoxicity and PTC readthrough ability in the different reporters based on the results presented in Figures 1 and 2. All cyclic peptides were encoded with a cysteine to facilitate cyclization (34). Remarkably, eight of the ten readthrough-promoting cyclic peptides contained a tryptophan residue at the first randomized position (Table 1). This feature is likely to support their nonsense-suppression activity, since tryptophan is not enriched at this specific position in the total cyclic peptide library (38). Exceptions to this are CP55, which contains another aromatic amino acid, tyrosine, at the same position, and CP35, which generally differs from the other cyclic peptides with a positively charged lysine and three hydrophobic isoleucines. Apart from a general preference for phenylalanine at the second position, the presence of a hydrophobic residue at the fifth randomized position also appeared to be favorable (Table 1). Altogether, the ten readthrough-stimulating cyclic peptides were verified to be expressed and processed in yeast, to hold the ability to suppress various nonsense mutations in reporter genes at differing efficiencies and specificities, and to contain structural similarities indicating the presence of functional sequence motifs.

**Table 1:**
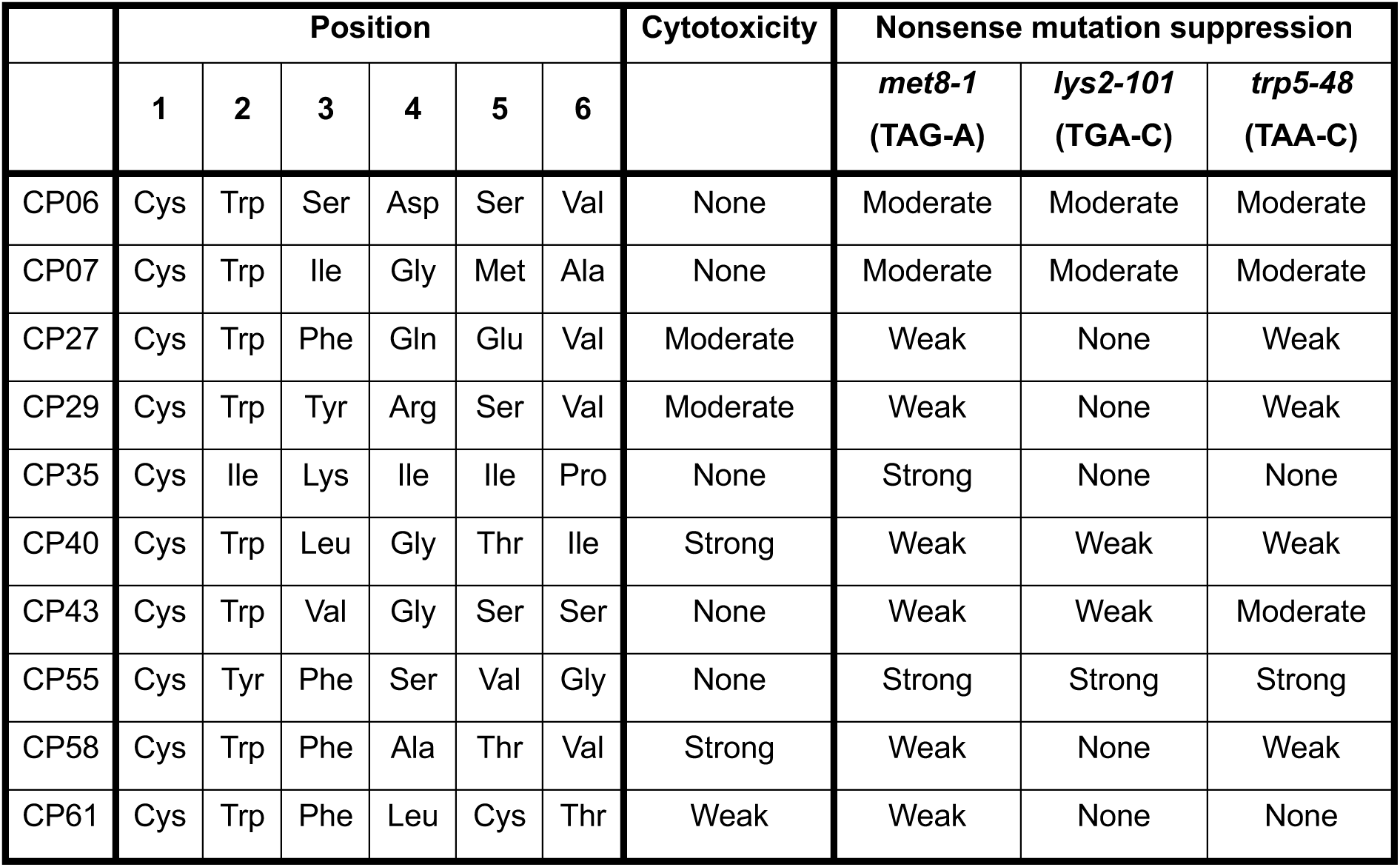
Sequences of ten selected readthrough-promoting cyclic peptides. Constructs encoding readthrough-promoting cyclic peptides (CPs) were subjected to Sanger sequencing to obtain the amino acid compositions of the cyclic peptides. Each cyclic peptide contains a constant cysteine followed by five amino acid positions that were randomized in the total DNA library. Note that since peptides are cyclized head-to-tail, position 6 is connected to the cysteine through the peptide backbone. The semi-qualitative assessment of cytotoxicity and nonsense mutation suppression ability is based on results shown in Figures 1 and 2.

### Four cyclic peptides display strong readthrough promotion and no cellular toxicity

To gain a quantitative assessment of how candidate cyclic peptides affect cell growth under non-selective and selective conditions, experiments were conducted using cells expressing cyclic peptides in liquid cultures. Four readthrough-promoting cyclic peptides, namely CP06, CP07, CP35, and CP55, were chosen as the lead candidates based on their strong readthrough promotion and lack of growth inhibition on solid medium (Table 1). Cell growth was monitored via the optical density at 600 nm (OD_600_) at specific time points.

General cell growth under conditions not selecting for readthrough promotion was monitored by culturing cells in a methionine-containing medium added galactose to induce cyclic peptide expression and omitted leucine to select for cells transformed with cyclic peptide-encoding constructs. Cultures were established using starter cultures containing galactose, thereby ensuring that the cells were already expressing the cyclic peptides from the beginning of the experiments. The relatively slow general growth of the yeast cultures was likely due to the use of galactose as the sole carbon source. Any potential cellular toxicity mediated by expression of the readthrough cyclic peptides was assessed by comparing growth to cultures expressing a random cyclic peptide as a negative control. Notably, almost no reduction in cell growth was observed for any of the cultures expressing the readthrough-promoting cyclic peptides (Figure 3A). At 24 and 48 hours, growth was even significantly enhanced for cultures expressing CP06, CP07, and CP35 compared to the control. Growth of the culture expressing CP55 was enhanced at 24 hours, but with a small, albeit significant, reduction in cell numbers (mean OD_600_ of 4.2 for CP55 and 4.4 for random cyclic peptide) at 48 hours (Figure 3A).

**Figure 3.**
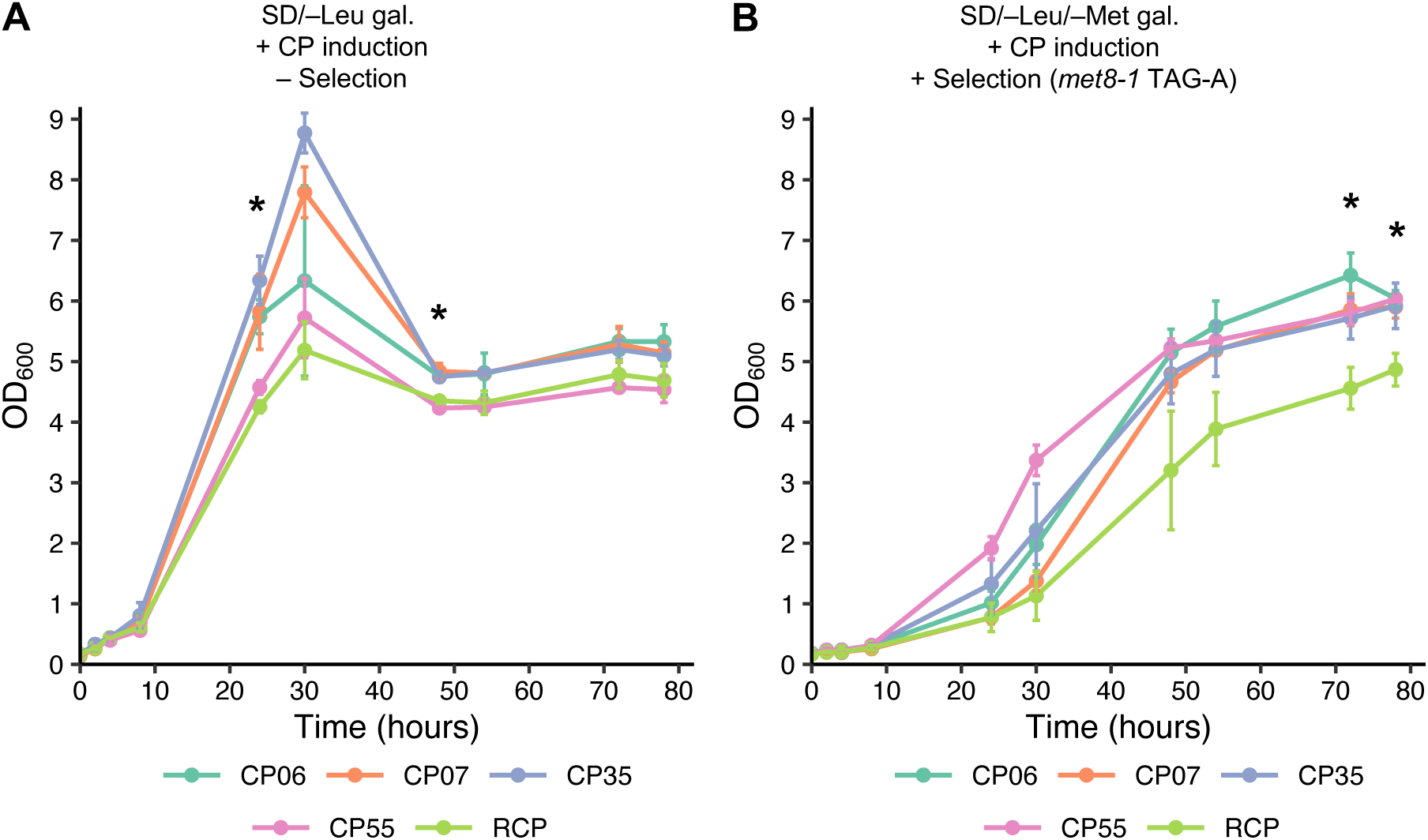
The most promising readthrough-enhancing cyclic peptides quantitatively show prominent readthrough and no cytotoxicity in *S. cerevisiae*. Constructs encoding readthrough-promoting cyclic peptides (CPs) were isolated and re-transformed into fresh *S. cerevisiae* IS110-18A cells, which were then grown in liquid culture. The presence of galactose (gal) induces CP expression, while the omission of leucine (–Leu) selects for transformed cells. The additional exclusion of methionine (–Met) allows for the selection of cells expressing CPs capable of suppressing a nonsense mutation present in the *met8* (*met8-1* TAG-A) gene. (**A**) Growth of cells under non-selective conditions. Potential cytotoxicity was assessed by comparison to cells expressing a random cyclic peptide (RCP). (**B**) Growth of cells under non-selective conditions. Growth enhancement through nonsense mutation suppression was assessed by comparison to cells expressing RCP. Shown are the mean ± SD of three biological replicates. *p<0.05 using Student’s t-test individually comparing readthrough-promoting CPs to RCP control, only marking time points where all CPs are significantly different from RCP.

Nonsense mutation suppression was assessed by additional omission of methionine and monitoring cell numbers in the same manner and timeframe as for the cultures without selection pressure. All cultures expressing readthrough-promoting cyclic peptides showed significant increases in cell numbers compared to control cultures, indicating nonsense-mutation suppression in the *met8-1* reporter gene (Figure 3B). The relatively high growth of control cultures can possibly be explained by individual cells within the cultures acquiring random genomic mutations over time that allow them to expand in the selective medium. Overall, the same trends were seen as for the assays on solid medium with CP55 being the superior nonsense-suppressing candidate, showing significantly enhanced cell growth from 24 hours of culturing and onwards, followed by CP35 (Figures 1C and 3B). Interestingly, while growth enhancement for CP06 and CP07 seemed identical in spot assays (Figure 1C), the application of OD_600_ measurements to assess cell growth revealed CP06 to be better than CP07 in promoting stop-codon readthrough (Figure 3B). In brief, the four most promising readthrough-promoting cyclic peptides display no observable cellular toxicity in yeast and support robust nonsense mutation suppression.

### Cyclization and most amino acid positions of CP55 are important for readthrough

Mutational analysis, including alanine scanning, was conducted on the lead readthrough-promoting candidate, CP55 (Figure 4A), to gain a better understanding of structural motifs contributing to its functionality. Mutagenesis was done at the DNA level by substituting codons in the cyclic peptide-encoding DNA construct. C1 was mutated to serine (instead of alanine) to investigate the importance of a thiol group, while still retaining the ability of the peptide to cyclize (60). The remaining residues were individually mutated to alanine. Y2 was also mutated to tryptophan to investigate if tryptophan is favorable in the context of CP55’s sequence since most other identified readthrough cyclic peptides have a tryptophan succeeding cysteine (Table 1). Moreover, constructs encoding non-mutated CP55 with point mutations in the intein gene segments were made to assess, if the readthrough-promoting activity is facilitated by the cyclized peptide, or by an unprocessed or partially processed intein-peptide intermediate (41). The introduction of the mutations T69A and H72A in the intein N segment blocks the first step in intein processing (43), while the mutations H24L and F26A in intein C inhibit the final processing step that leads to the release of cyclized peptides (44).

**Figure 4.**
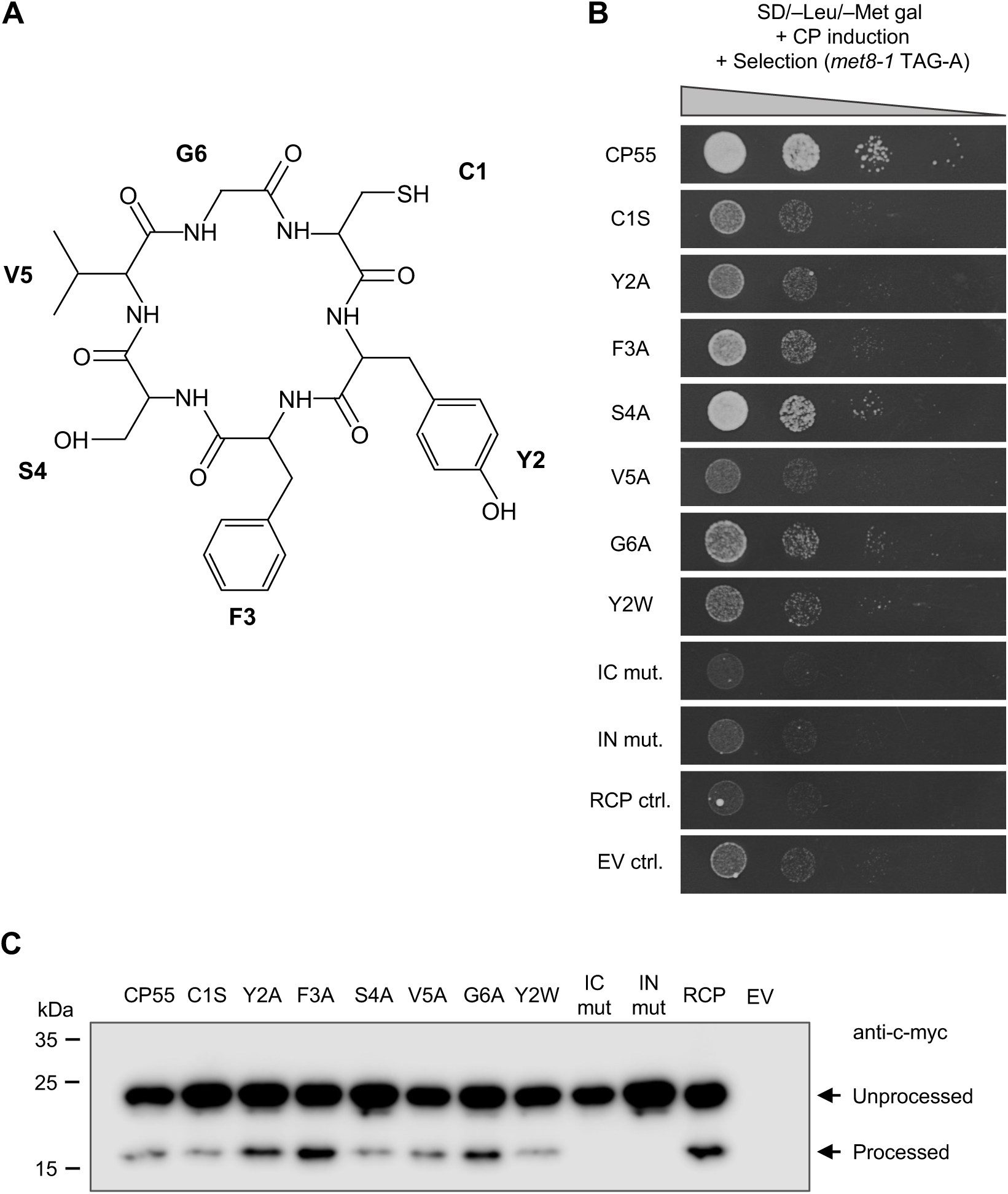
Mutational analysis of CP55 shows that most amino-acid side chains contribute to its nonsense-mutation suppression activity. (**A**) Chemical structure of the readthrough cyclic peptide (CP) CP55, containing L-amino acids. The six mutated residues are indicated. (**B**) Constructs encoding originally selected CP55 and mutants thereof were transformed into *S. cerevisiae* IS110-18A and spotted in serial ten-fold dilutions on the selective medium. The presence of galactose (gal) induces CP expression, while the omission of leucine (–Leu) selects for transformed cells. The additional omission of methionine (–Met) selects for cells expressing CPs capable of suppressing a nonsense mutation present in the *met8* (*met8-1* TAG-A) gene. Growth enhancement through nonsense-mutation suppression was evaluated by comparison to cells expressing a random cyclic peptide (RCP) or an empty vector (EV), encoding intein segments without insertion of randomized sequences resulting in a frameshift. Images were taken after four days of incubation and are representative of three independent experiments. (**C**) Western blotting detects the expression and processing of CP55 and its mutant derivatives by means of an anti-c-myc antibody. Lysates of *S. cerevisiae* IS110-18A cells transformed with the relevant CP55-encoding constructs were analyzed, along with cells expressing EV or RCP as controls. IC, intein C; IN, intein N.

Constructs encoding CP55 and its mutated variants were transformed into *S. cerevisiae* IS110-18A cells, cultured, and spotted on a selective medium lacking methionine. Control plates without selection pressure demonstrated that none of the CP55 mutants had any effect on general cell growth (Figure S5). Strikingly, almost all mutations either abolished or greatly reduced the nonsense-suppression activity of CP55 (Figure 4B). S4 was the only residue that did not seem to contribute substantially to the readthrough-promoting activity, as mutating this amino acid to alanine only resulted in a minor growth reduction as compared to non-mutated CP55. Importantly, mutations in the intein segments that hinder processing completely abolished the activity, demonstrating that the active readthrough-stimulating compound is the fully cyclized CP55 that results after splicing and release from the intein domains (Figure 4B).

Western blotting confirmed that all variants of CP55 were highly expressed, and all displayed processing, including the CP55 C1S mutant (Figure 4C; unprocessed blot Figure S3B). This suggests that the reduced nonsense-mutation suppression of the CP55 mutants is not due to deficits in expression or cyclization. Furthermore, the lack of processing was confirmed for the intein mutants (Figure 4C). To summarize, mutational analysis demonstrated that most of the amino acid side chains in CP55 contribute to its readthrough activity. The results also showed that the cyclized peptide, and not an intein-peptide intermediate, is the active readthrough-enhancing molecule.

### CP55 promotes readthrough by direct interactions with the protein synthesis machinery

In this study, several cyclic peptides have been identified that promote cell survival in *S. cerevisiae* as a consequence of translational readthrough in essential biosynthesis genes (Table 1). Their selection was not based on a specific mechanism of action, and multiple distinct cellular targets could lead to this observed readthrough. Many readthrough-promoting molecules have been demonstrated to function by a direct binding to the ribosome or other core translation factors (16,17,53,61). However, the modulation of other cellular pathways, ultimately leading to an increase in readthrough, has also been described, including interference with the NMD pathway (62) or alterations of tRNA-modifying enzymes (63).

To gain mechanistic insights into the observed readthrough-promoting activity of the identified cyclic peptides, several *in vitro* biochemical eukaryotic translation assays were employed. The assays were used to investigate CP55, which was the lead candidate displaying the strongest readthrough promotion in yeast with minimal toxicity (Figure 3), and where most of the amino acid side chains of the fully circularized peptide seem to be involved in target binding (Figure 4). The assays employ eukaryotic 80S ribosomes derived from brine shrimp (*A. salina*), programmed with mRNA sequences containing an IRES sequence to facilitate translation initiation in the absence of initiation factors. The assays also contain purified yeast or *E. coli* aminoacyl-tRNAs, yeast elongation factors, and human termination factors (see Materials and Methods for details). Through IRES-mediated initiation and elongation *in vitro*, ribosomes complexes are formed that contain Phe-Lys-Val-Arg-Gln-tRNA^Gln^ in the P site and a UGA stop codon in the A site (Figure 5A) (52,53). These complexes are termed Stop-POST5 which denotes that the ribosome is in its post-translocation state with a termination codon in the A site, and that the nascent peptide at this point contains five amino acids. The addition of a release factor complex (RFC) to the reaction results in translation termination and the release of the peptide (Figure 5A) (48,52). In contrast, adding elongation factor eEF1A·GTP and a near-cognate tRNA (Trp-tRNA^Trp^) fosters readthrough by accommodation of the tRNA into the A site containing the UGA termination codon. Following accommodation and peptide transfer, the A site contains Phe-Lys-Val-Arg-Gln-Trp-tRNA^Trp^ (PRE6), and subsequent translocation by elongation factor eEF2·GTP moves the hexapeptidyl-tRNA to the P site (to produce POST6) (Figure 5A) (16,53). As a control, the experiments also included the Trp-POST5 complex which contains a cognate TGG codon encoding tryptophan in the A site. For the experiments, CP55 was synthesized chemically and cyclized head-to-tail (see Materials and Methods). The identity of the cyclic peptide was verified by mass spectrometry.

**Figure 5.**
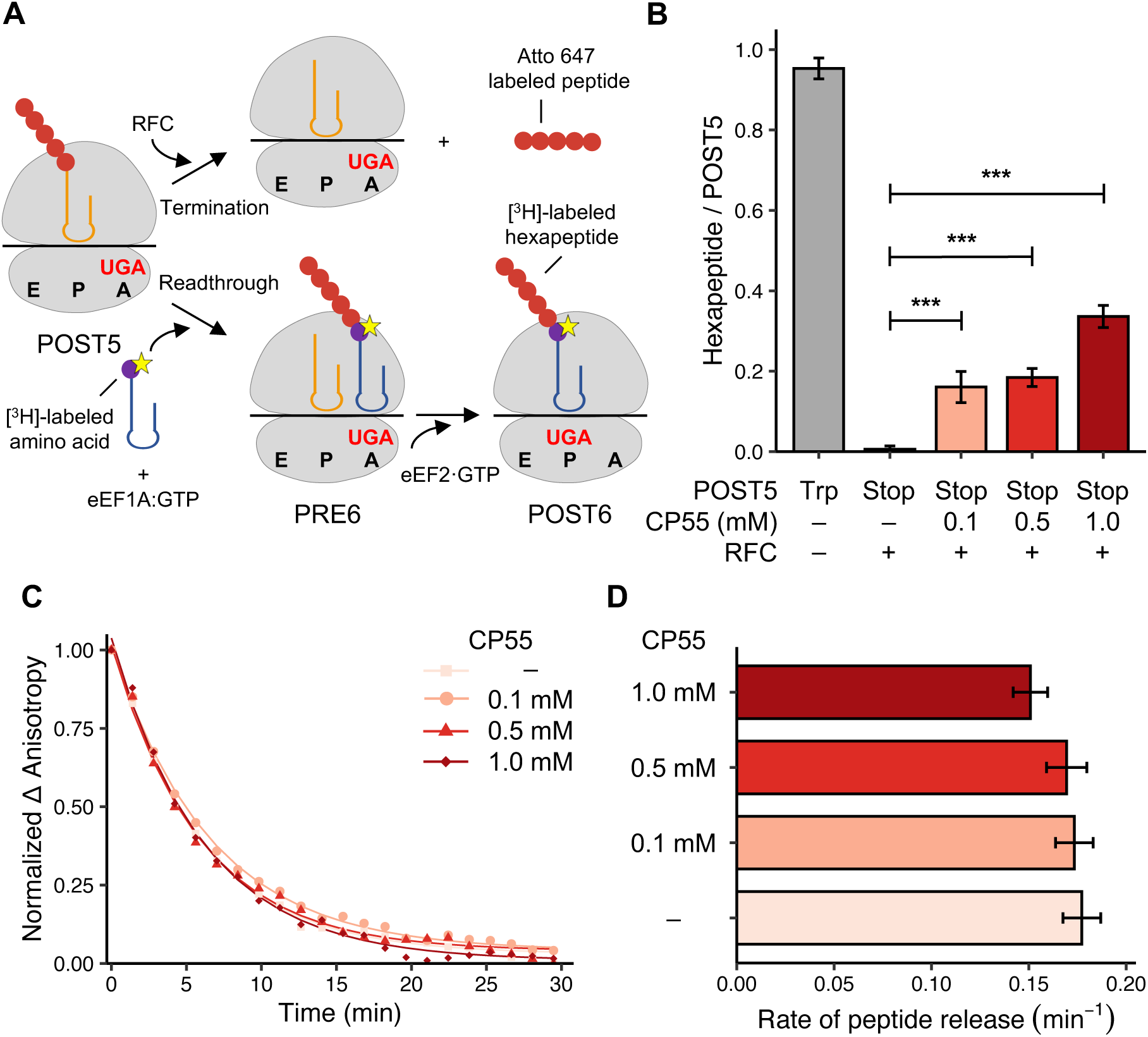
*In vitro* translation assays suggest that CP55 promotes readthrough via interference with decoding. (**A**) Schematic of applied termination and readthrough assays. The starting point for all assays is the POST5 complex denoting a ribosome with a UGA termination codon in the A site and a pentapeptidyl-tRNA in the P site. In termination assays (top), a release factor complex (RFC) facilitates translation termination, which can be monitored through the release of an Atto 647-labeled peptide. In readthrough assays (bottom), the addition of a near-cognate tRNA and eukaryotic elongation factor eEF1A results in tRNA accommodation and peptide-bond formation (forming PRE6), and the presence of eEF2 facilitates subsequent translocation (forming POST6). For co-sedimentation assays, readthrough is monitored through the insertion of a [^3^H]-labeled amino acid that co-sediments with the POST6 complex. (**B**) Co-sedimentation assay measuring readthrough by the amount of formed hexapeptide relative to POST5 at various concentrations of cyclic peptide (CP) CP55. A control reaction was included using a POST5 complex with a UGG (Trp) codon in the ribosomal A site, to measure cognate amino-acid insertion. The experiment was performed in the presence of RFC. Shown is the mean ± SD of duplicate reactions from two independent experiments. ***p<0.001 using one-way ANOVA. (C, D) Termination assay measuring rates of peptide release via changes in fluorescence anisotropy (Δ Anisotropy) resulting from the release of Atto 647-labeled nascent peptides after peptidyl-tRNA hydrolysis. Experiments were conducted in the presence of varying concentrations of CP55 as indicated. (**C**) Normalized 1′ Anisotropy as a function of time. Traces were fit using a one-phase exponential decay model. (**D**) Rates of peptide release from the fitted curves shown in (C). Shown is the mean ± SD of three independent experiments.

First, a potential *in vitro* effect of CP55 was investigated in a co-sedimentation readthrough assay, where the incoming near-cognate tRNA is radioactively labeled (Figure 5A). Upon near-cognate tRNA binding and amino acid insertion, a [^3^H]-labeled hexapeptide is formed, which co-sediments with ribosomes upon ultracentrifugation (Figure 5A). Measuring the radioactivity of resuspended ribosomes relative to the amount of initial POST5 complexes quantifies readthrough (16,53). The experiment was performed in the presence of RFC, which creates competition in the A site between RFC and near-cognate tRNA binding. Strikingly, CP55 increased readthrough in a concentration-dependent manner, indicating that CP55 interacts directly with the ribosome or core translation factors (Figure 5B). Of note, almost no readthrough was observed in the absence of CP55 due to the addition of RFC (Figure 5B) (16), suggesting a prominent readthrough effect of CP55.

When inhibiting release-factor activity, translation termination is impaired, which can indirectly result in increased readthrough due to the decreased occupancy of RFC in the ribosomal A site (Figure 5A) (16). Indeed, the readthrough-promoting molecule ataluren has been shown to promote readthrough by inhibiting eRF1/eRF3-mediated peptidyl-tRNA hydrolysis (16,52). To investigate if CP55 affects the termination process directly, the cyclic peptide was tested in a termination assay employing POST5 complexes in which the pentapeptide in the P site was fluorescently labeled with Atto 647. Upon termination and release of the labeled peptide, termination was quantified by changes in Atto 647 fluorescence anisotropy (Figure 5C) (52). Rates of peptide release were estimated from the resulting curve fits (Figure 5D). CP55 did not affect the rate of translation termination, except for a slight decrease at a high concentration (Figure 5D). This indicates that CP55 promotes readthrough by interfering directly with ribosomal decoding in a mechanism similar to that of aminoglycoside antibiotics (16). Indeed, performing the readthrough co-sedimentation assay in the absence of release factors retained a readthrough-stimulating effect of CP55, supporting this hypothesis (Figure S6).

To further substantiate the mechanism behind the readthrough-promoting effect of CP55 *in vitro*, the cyclic peptide was tested in single-molecule FRET by applying TIRF microscopy (53,54). The experimental set-up employed a biotinylated Stop-POST5 complex containing a Cy5-conjugated peptidyl-tRNA^Gln^ in the P site bound to the microscope slide. Readthrough was determined by the binding of a near-cognate Cy3-conjugated Trp-tRNA^Trp^ in the A site, producing a tRNA-tRNA FRET signal between the two fluorophores with a FRET efficiency of ∼0.5 (Figure S7). Formation of PRE6 complexes (with tRNAs occupying both the P- and A-sites), as detected by colocalizing FRET donor and acceptor signals on the microscope slides to identify FRET events, was prominent for Trp-POST5 complexes (Figure 6A) but substantially less for Stop-POST5 complexes (Figure 6B). The addition of CP55 to the Stop-POST5 complexes increased the number of PRE6 complexes formed, supporting the readthrough-promoting effect of CP55 (Figure 6C). Quantification of readthrough by the number of PRE6 complexes formed displayed a concentration-dependent increase upon the addition of CP55 (Figure 6D). Together, these results strongly support that CP55 can promote readthrough by modulating ribosomal decoding directly in eukaryotic cells.

**Figure 6.**
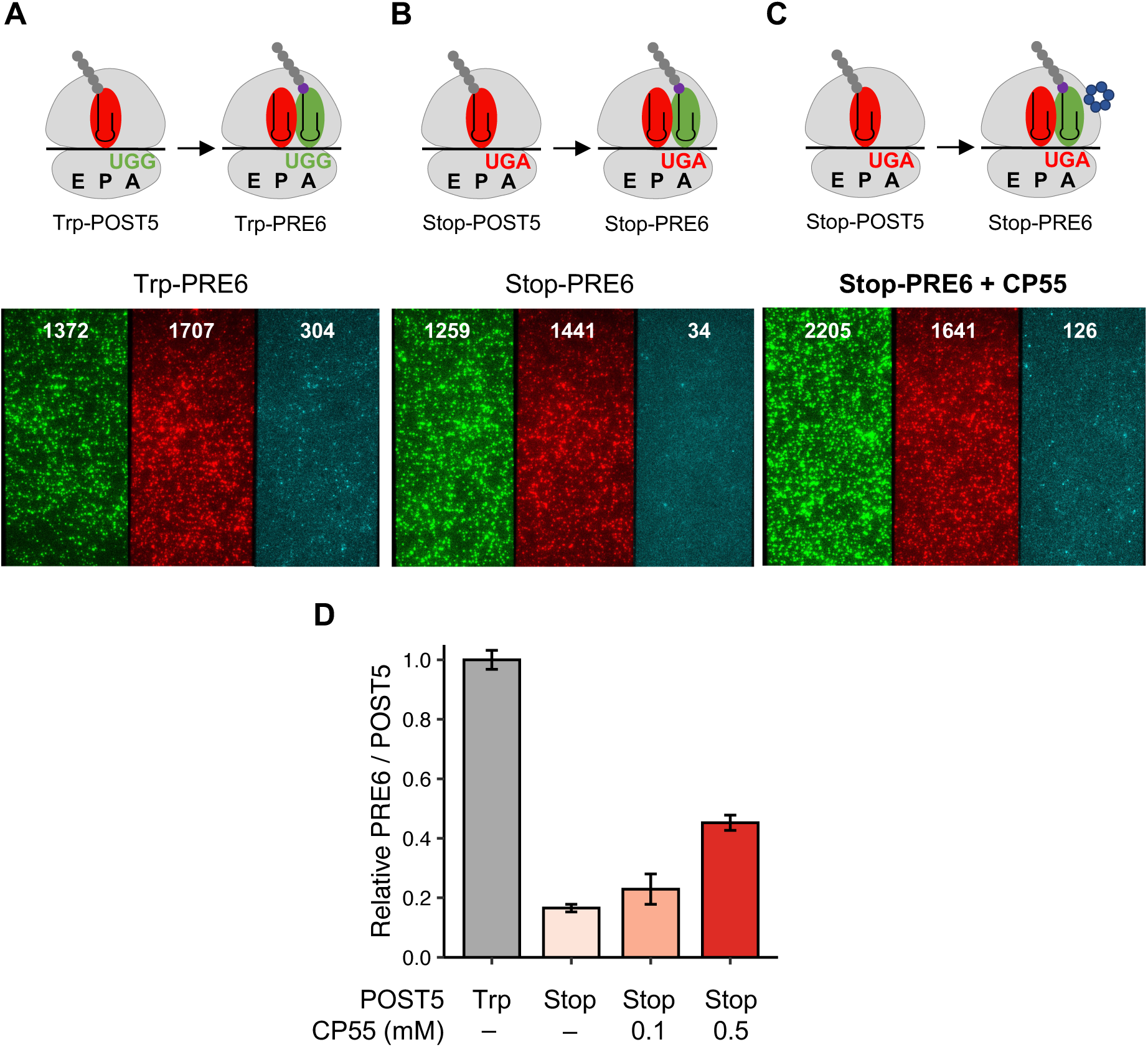
Single-molecule FRET supports a readthrough-enhancing effect of CP55. (**A-C**; upper part) Readthrough is quantified as the fraction of PRE6 complexes formed relative to initial POST5 complexes, as measured by FRET between two fluorescently labeled tRNAs using TIRF microscopy upon alternating laser excitation. (**A-C**; lower part) Signals from the Cy3 donor (left), the Cy5 acceptor (middle), and FRET signals (detected upon colocalization of the two first panels; right) of PRE6 complexes formed from ribosomes containing either (**A**) a cognate UGG (Trp) codon (Trp-PRE6) or a UGA termination codon (Stop-PRE6) in the (**B**) absence or (**C**) presence of cyclic peptide (CP) CP55. Representative images are shown. White numbers on panels indicate the number of Cy3 (green), Cy5 (red), and FRET (cyan) particles on the imaged microscope slide. (**D**) Quantification of the fraction of FRET signals, normalized to the amount of formed Trp-PRE6, with increasing concentrations of CP55. The number of experiments ranged from 2–4 for each condition. Shown is the mean ± SEM which is calculated from the total number of spots.

## DISCUSSION

Despite many years of pre-clinical and clinical studies (14,15), ataluren remains the only clinically approved readthrough-promoting drug (26). However, ataluren only benefits a small subset of patients suffering from Duchenne muscular dystrophy (25,26), indicating a need for new strategies to promote readthrough at PTCs. In the present study, we have identified ten novel cyclic peptides capable of suppressing genomic nonsense mutations in *S. cerevisiae* (Table 1), with four showing no detectable cellular toxicity (Figure 3A). Further work on the best performing candidate, CP55, revealed that most residues in the cyclic peptide engage in target binding (Figure 4) and that CP55 promotes readthrough by binding the eukaryotic translation machinery and interfering with ribosomal decoding of a stop codon (Figures 5 and 6). We reasoned that cyclic peptides, being larger than current small-molecule drugs targeting nonsense mutations, would be superior in inhibiting or modulating protein-protein or protein-RNA interactions that promote PTC readthrough. This mechanism could plausibly be in play for the observed readthrough effect of CP55. To our knowledge, the naturally derived antimicrobial compound negamycin is the only peptide-derived molecule reported to suppress nonsense mutations (64,65). Negamycin is a linear, modified dipeptide (66) and bears limited structural resemblance to our identified readthrough-stimulating cyclic peptides (Figure 4A).

The cyclic peptides facilitating readthrough of PTCs were discovered from a yeast SICLOPPS library (38) employing a selection strategy that couples cell survival and growth to nonsense mutation suppression (Figure 1). The SICLOPPS system allows for intracellular expression of circularized peptides, thereby making it possible to perform *in vivo* selection of cyclic peptides based on functionality and absence of cytotoxicity. This strategy is advantageous relative to *in vitro* affinity-based screening methods since strong binding affinity does not necessarily result in modulation of cellular function (29). SICLOPPS has mostly been applied for screens in bacterial cells, although eukaryotic protein interactions can also be assayed in bacteria via heterologous expression (67,68). Moreover, SICLOPPS libraries have mainly been subject to selection via bacterial two-hybrid systems, where cyclic peptide inhibitors of specified protein-protein interactions are identified (37,67,68). Only two studies have reported the use of SICLOPPS in eukaryotic cells to identify cyclic peptides promoting a certain phenotype, without specifying an exact target as is the case for two-hybrid systems (41,69). Our approach arguably makes determining the mechanism of action for the identified cyclic peptides a challenge, but it also allows for an unbiased identification of compounds modulating a specific pathway. Using this strategy, we discovered cyclic peptide suppressors of nonsense mutations, possibly with different cellular targets.

A subset of the identified readthrough-promoting cyclic peptides caused growth inhibition when expressed in yeast (Table 1). In the applied selection system, nonsense-mutation suppression is coupled to cell growth. Thus, it cannot be ruled out that very strong readthrough-inducing cyclic peptides are simultaneously growth-inhibitory, thus masking their stimulation of readthrough due to a reduction in cell viability. Of note, PTCs are generally more prone to readthrough compared to natural termination codons, ensuring that drug-induced readthrough mainly targets PTCs (58,70,71). Yet, the observed reductions in cell growth (Figure 1E) could potentially be due to potent readthrough induction targeting both PTCs and natural termination codons. Compounds with high cytotoxicity would not be relevant as potential therapeutics. For example, aminoglycoside antibiotics are strong nonsense mutation suppressors (72,73), but cannot be used at therapeutic doses due to their toxicity (74). Our selection system addresses this problem by linking nonsense mutation suppression to cell survival, ensuring that the identified cyclic peptides do not have any major cytotoxic effects. Importantly, four of the identified readthrough cyclic peptides did not affect yeast cell viability (Figure 3A).

It is well established that the specific termination codon and surrounding sequence affect both basal and induced readthrough levels (11,58,75–77). The UGA stop codon is permissive to readthrough, whereas the UAA stop codon is a strong signal for translation termination and therefore a difficult target for readthrough promotion (11). Moreover, a purine following the termination codon also decreases the likelihood of readthrough (6,58). In the current study, almost no nonsense-mutation suppression was observed for the TAA nonsense mutations present in *ura4* and *ilv1* either by the identified readthrough-promoting cyclic peptides (Figure S4 A and C) nor by G418 (Figure S4B). Whereas the high background observed for *lys2-101* (TGA-C) (Figure 2C) could potentially be explained by UGA termination codons permitting high levels of basal readthrough (11,58), it is currently unknown why such a high background growth is observed upon selection for suppression of the *trp5-48* (TAA-C) nonsense mutation. Regardless, the most promising candidates CP06, CP07, and CP55 were able to suppress nonsense mutations of all three termination codons (Figures 1C, 2C, and 2D). One exception was CP35, which seemed to carry a high specificity for UAG PTCs (Table 1). This sequence specificity of CP35 warrants further investigation.

The readthrough-enhancing cyclic peptides reported here were identified in a selection system using *S. cerevisiae* as a model organism. However, our ultimate goal is to identify nonsense-mutation suppressors with the potential of being developed into therapeutics to treat human diseases. Yeast was chosen as a eukaryotic cell model of translation termination due to well-established methodologies of linking growth enhancement to the suppression of various genomic mutant alleles, and the ability to achieve a high number of transformed cells when introducing a DNA-encoded library. In addition to suppressing nonsense mutations in *S. cerevisiae*, CP55 was shown to promote readthrough *in vitro* using ribosomes derived from shrimp cysts, suggesting possible capacity of CP55 in suppressing PTCs in human cells, due to the high conservation of the core translation machinery across eukaryotic species (6,17). Notably, CP55 displays characteristics similar to aminoglycosides in the applied *in vitro* assays, where readthrough promotion is seen even in the absence of release factors with no apparent effects on translation termination (Figures 5B, 5D, S6, and 6D) (16,53). Structural studies have revealed a single tight binding site of aminoglycosides in the decoding center of *S. cerevisiae* ribosomes (17,78). Moreover, many studies have demonstrated a readthrough-stimulating effect of aminoglycoside antibiotics in both yeast and human cells (40,79). It is feasible that CP55 promotes readthrough by binding to, or in the vicinity of, the ribosomal A site. We are currently investigating if the identified readthrough-promoting cyclic peptides are active in human cells.

The use of SICLOPPS libraries and relevant library selection protocols is a powerful tool for the *de novo* discovery of cyclic peptides with the ability to bind and modulate difficult cellular targets (30,31). Our constructed library encodes circularized peptides of six amino acids (38). Keeping the size of the identified cyclic peptides small likely results in an increased cell permeability (27,80) and makes further development into a therapeutic drug more accessible. Of note, studies have reported on successful chemical modification of SICLOPPS-derived small, cyclic peptides with the scope of developing the initial compounds into more drug-like molecules with increased potency and improved pharmacological properties (30,81).

In summary, we report the fruitful selection of ten novel cyclic peptides from a SICLOPPS library that can suppress nonsense mutations in *S. cerevisiae*. The four most promising candidates are non-cytotoxic and can promote readthrough at PTCs in various genes and sequence contexts. Our lead candidate, CP55, was shown to interact directly with the eukaryotic protein synthesis machinery and promote readthrough by interference with ribosomal decoding without affecting the termination process. Further studies and developments will hopefully expand this work into cyclic peptides functioning in human cells, and thus carry the potential to be used in treating diseases that are the result of nonsense mutations.

## SUPPLEMENTARY DATA

A single file containing Table S1-S2 and Figures S1-S7, with accompanying legends.

## ACKNOWLEDGEMENTS

We thank Dr. Alireza Baradaran-Heravi (University of British Columbia) for sharing yeast strains and Prof. Ali Tavassoli (University of Southampton) for sharing a bacterial SICLOPPS library. We acknowledge Anders Løbner-Olesen (University of Copenhagen) and Christian Kroun Damgaard (Aarhus University) for valuable feedback. We thank Manfred Schmid for yeast expression vectors. Bente Flügel Majgren is acknowledged for technical assistance.

## FUNDING

This work was supported by a grant from the Novo Nordisk Foundation (to C.R.K.); Tømrermester Jørgen Holm og Hustru Elisa f. Hansens Mindelegat (to C.R.K.); Fabrikant Einar Willumsens Mindelegat (to C.R.K.); and NIH grant R35GM118139 (to Y.E.G).

## Conflict of interest statement

A university-owned patent application (with C.R.K. and N.B. as inventors) of the identified readthrough-promoting cyclic peptides has been submitted.

