## Supplementary data for "Suppression of nonsense mutations by small, cyclic peptides"

| **Medium** | **Days of incubation before imaging** | |
|---|---|---|
| | Glucose | Galactose |
| **– Selection**<br>**SD/–Leu** | 2 | 3 |
| **+ Selection**<br>**SD/–Leu/–(Met/Lys/Trp/Ura/Ile)** | 3 | 4/5 |

**Table S1:** Days of incubation for imaging comparable yeast cell growth in spot assays. The presence of galactose in the medium induces cyclic peptide expression. The omission of leucine (–Leu) selects for transformed cells, while the omission of methionine, lysine, tryptophan, uracil, or isoleucine (–(Met/Lys/Trp/Ura/Ile)) selects for cells expressing cyclic peptides capable of suppressing a nonsense mutation present in their respective biosynthesis genes. As cells generally grow more slowly on galactose-containing medium compared to medium with glucose, representative images were taken after one additional day of growth on galactose-containing medium. Similarly, growth on selective medium was imaged after one or two additional days of growth to better capture the relatively slow growth-inducing effects of the identified cyclic peptides.

| Primer | Sequence 5'-3' | Purpose |
|---|---|---|
| GAL1_fwd | ATTTTCGGTTTGTATTACTTC | Sanger sequencing primer |
| GAL1_rev | GTTCTTAATACTAACATAACT | Sanger sequencing primer |
| CP55_C1S_1 | CGATCGCCCACAATTCCTACTTCTCGGTGG | Mutagenesis primer, C1S |
| CP55_C1S_2 | CCACCGAGAAGTAGGAATTGTGGGCGATCG | Mutagenesis primer, C1S |
| CP55_Y2A_1 | GATCGCCCACAATTGCGCCTTCTCGGTGGGCTGC | Mutagenesis primer, Y2A |
| CP55_Y2A_2 | GCAGCCCACCGAGAAGGCGCAATTGTGGGCGATC | Mutagenesis primer, Y2A |
| CP55_F3A_1 | GCCCACAATTGCTACGCCTCGGTGGGCTGCCT | Mutagenesis primer, F3A |
| CP55_F3A_2 | AGGCAGCCCACCGAGGCGTAGCAATTGTGGGC | Mutagenesis primer, F3A |
| CP55_S4A_1 | CCCACAATTGCTACTTCGCGGTGGGCTGCCT | Mutagenesis primer, S4A |
| CP55_S4A_2 | AGGCAGCCCACCGCGAAGTAGCAATTGTGGG | Mutagenesis primer, S4A |
| CP55_V5A_1 | CAATTGCTACTTCTCGGCGGGCTGCCTCAGTTTTG | Mutagenesis primer, V5A |
| CP55_V5A_2 | CAAAACTGAGGCAGCCCGCCGAGAAGTAGCAATTG | Mutagenesis primer, V5A |
| CP55_G6A_1 | GCTACTTCTCGGTGGCCTGCCTCAGTTTTGG | Mutagenesis primer, G6A |
| CP55_G6A_2 | CCAAAACTGAGGCAGGCCACCGAGAAGTAGC | Mutagenesis primer, G6A |
| CP55_Y2W_1 | GCAGCCCACCGAGAACCAGCAATTGTGGGCGA | Mutagenesis primer, Y2W |
| CP55_Y2W_2 | TCGCCCACAATTGCTGGTTCTCGGTGGGCTGC | Mutagenesis primer, Y2W |
| ICmut_1 | AAGACCTTAATGCTCTGCTAGCCAATGGGGCGATCG | Mutagenesis primer, H24L+F26A in intein C |
| ICmut_2 | AGCAGAGCATTAAGGTCTTGGGGAAGACCAATATCAAATATTCTTTGC | Mutagenesis primer, H24L+F26A in intein C |
| INmut_1 | GCTGCCTCTGACGCCCGCTTTTTAACCACCGATTATCAACTGTTG | Mutagenesis primer, T69A+H72A in intein N |
| INmut_2 | GGCGTCAGAGGCAGCTCGGATTACTGAGCCATCTTCCAATTCATATTC | Mutagenesis primer, T69A+H72A in intein N |

**Table S2:** Sequences and purpose of primers used in this study.

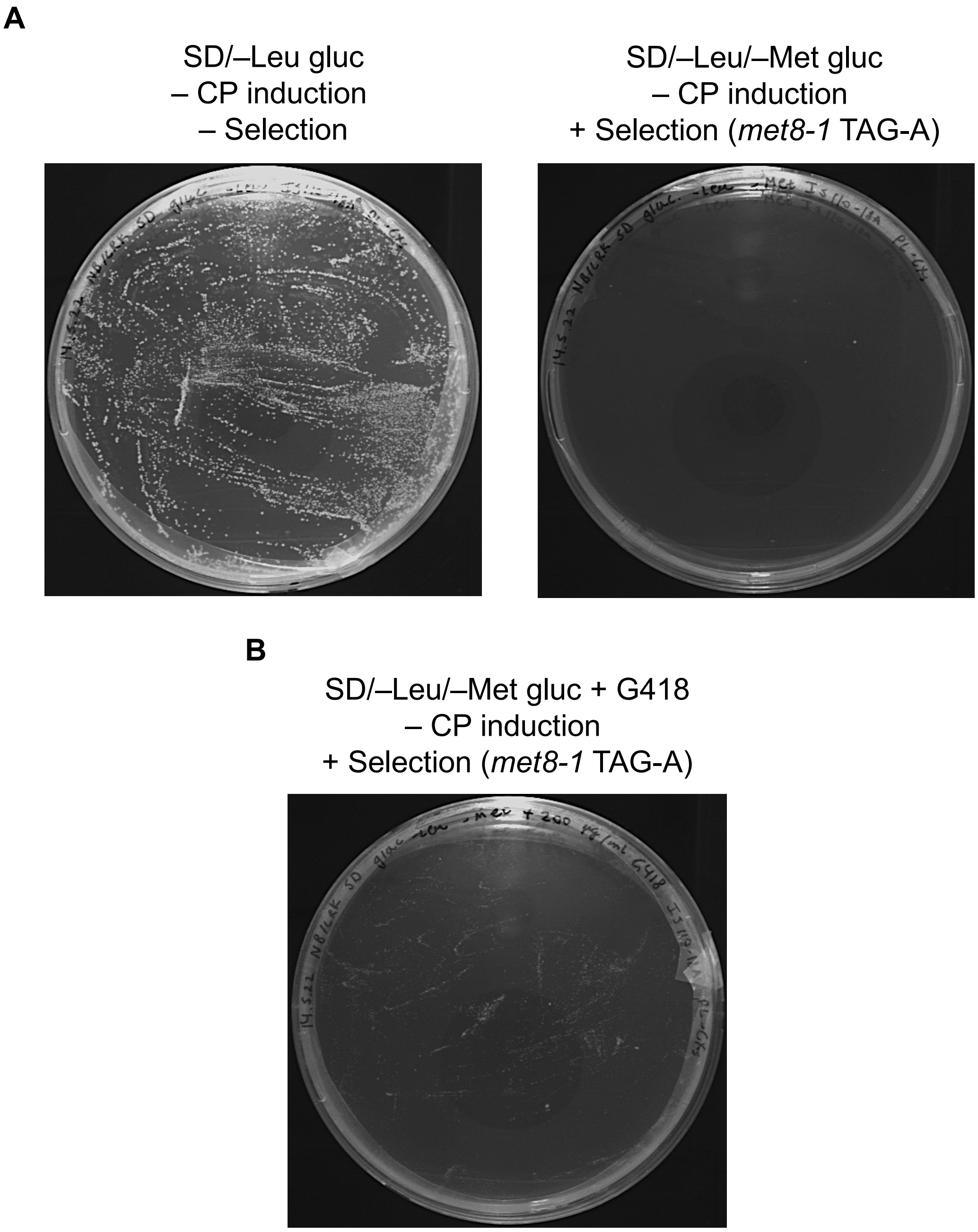


**Figure S1.** Control agar plates for the selection experiment without expression of cyclic peptides. Cyclic peptides (CPs) are not expressed when *S. cerevisiae* IS110-18A cells are grown in the presence of glucose (gluc). Omission of leucine (–Leu) selects for transformed cells. The additional exclusion of methionine (–Met) selects for nonsense-mutation suppression in the *met8* (*met8-1* TAG-A) gene. (**A**) Control plates with no induction of the SICLOPPS library, with (right) and without (left) selective pressure for readthrough of the nonsense mutation in *met8*. Images were taken after two days (left) or three days (right) of incubation. (**B**) Control plate with no induction of the SICLOPPS library, but with selection for nonsense-mutation suppression in the presence of the readthrough-inducing drug G418. The image was taken after three days of incubation.

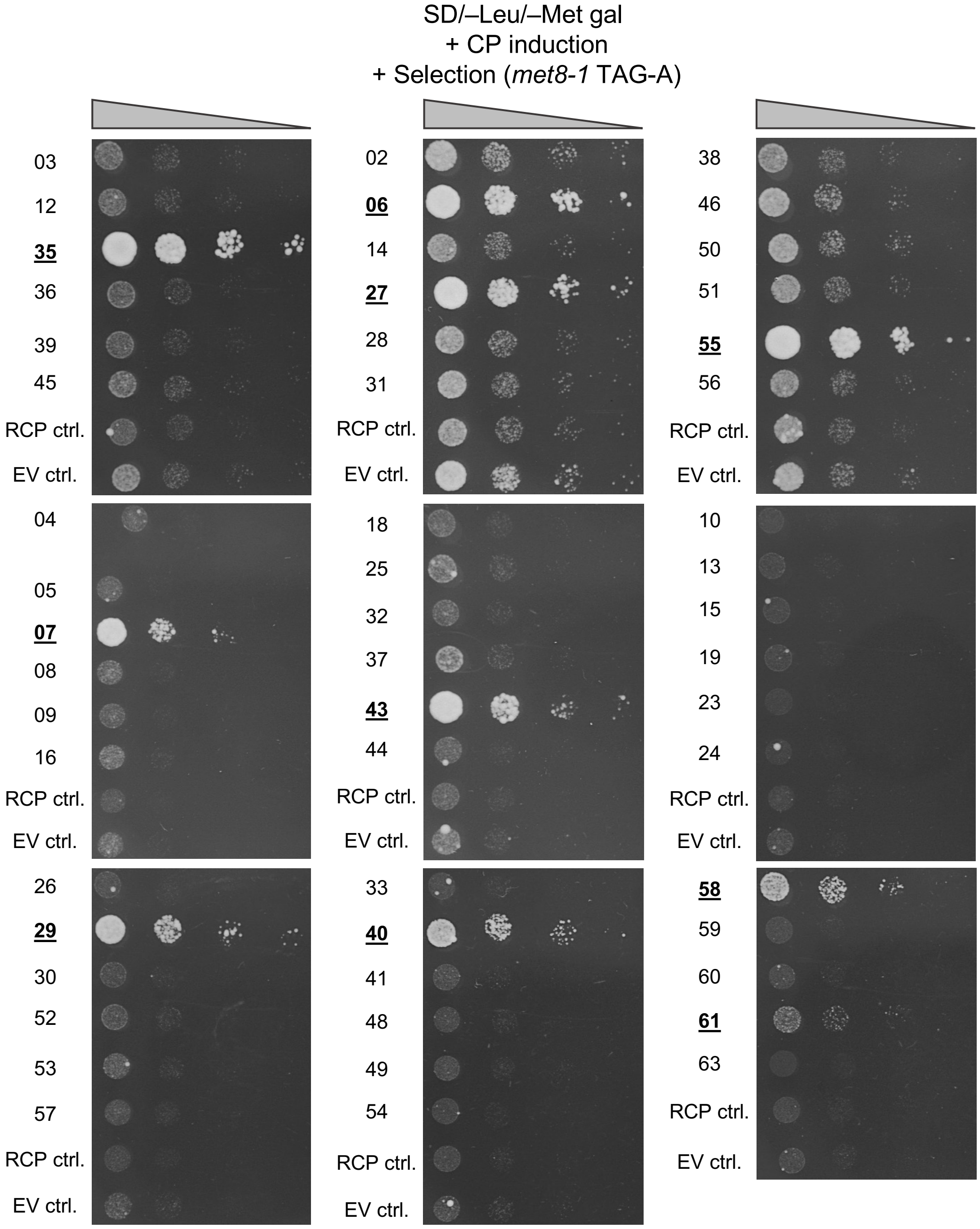


**Figure S2.** Validation of the readthrough-inducing potential of all selected cyclic peptides. Constructs encoding cyclic peptides (CPs) with readthrough-promoting potential were isolated from surviving colonies detected in the selection experiment (see example in Figure 1B). The constructs were re-transformed into fresh *S. cerevisiae* IS110-18A cells and spotted in serial ten-fold dilutions on the selective medium. The presence of galactose (gal) induces CP expression, while the omission of leucine (–Leu) selects for transformed cells. The additional omission of methionine (–Met) selects for cells expressing CPs capable of suppressing a nonsense mutation present in the *met8* (*met8-1* TAG-A) gene. Growth enhancement through nonsense-mutation suppression was evaluated by comparison to cells expressing a random cyclic peptide (RCP) or an empty vector (EV). CPs validated to suppress nonsense mutations when expressed in the yeast reporter strain are marked in bold and underlined. Images were taken after four days and are representative of two independent experiments.

**A**

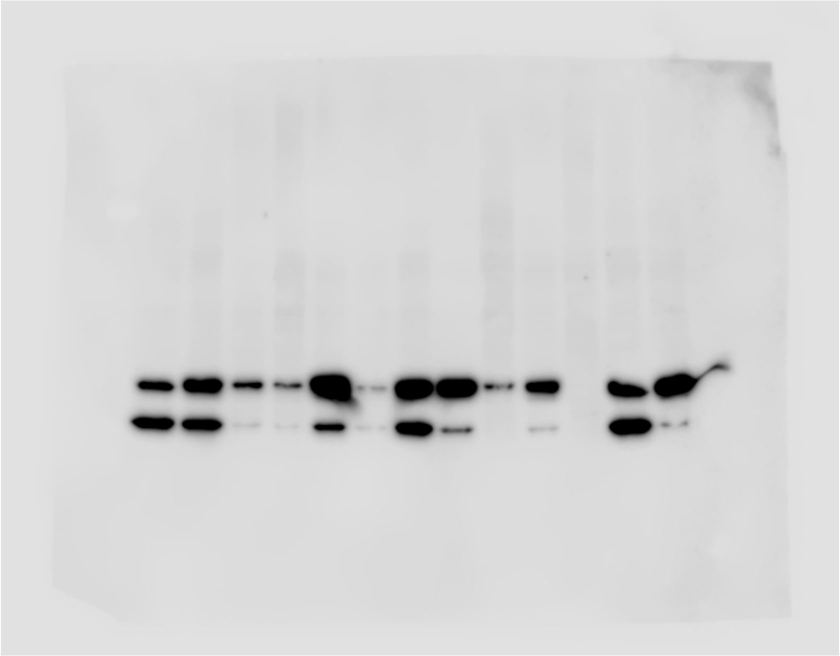

**B**

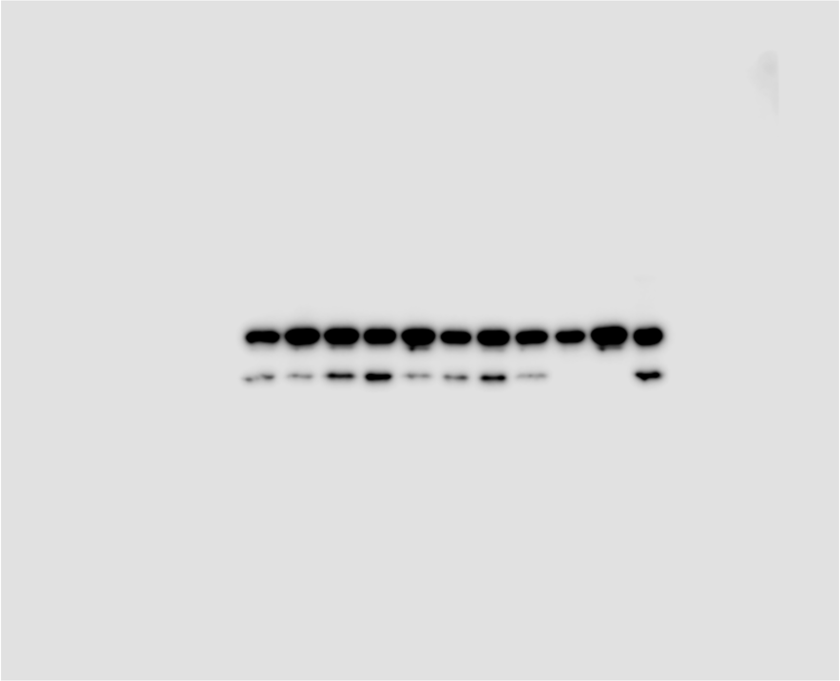

**Figure S3.** Unprocessed Western blot membranes. These blots are uncropped and unedited versions of blots shown in Figure 2B (**A**) and Figure 4C (**B**).

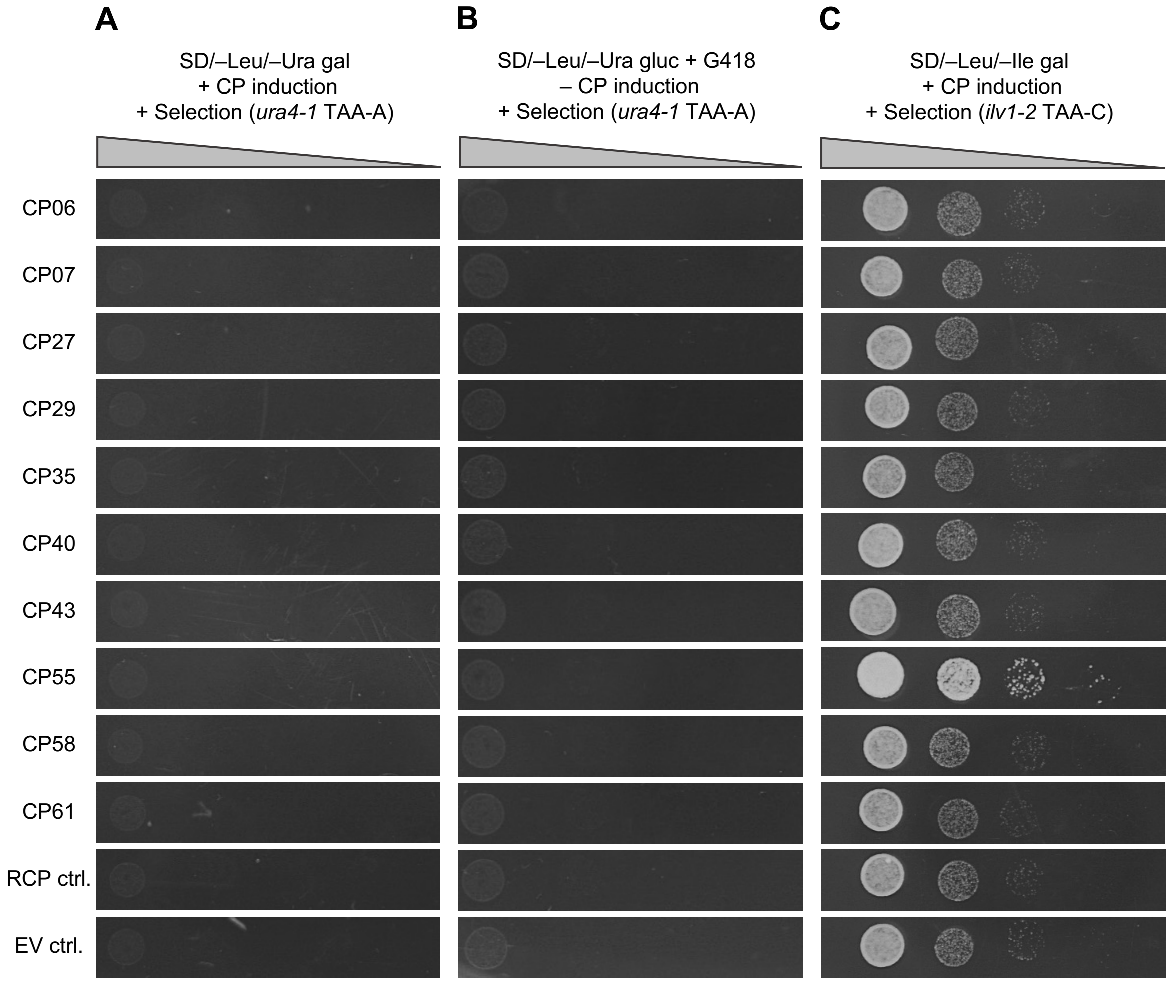


**Figure S4.** Selected, readthrough-promoting cyclic peptides showing no nonsense-mutation suppression in *ura4* and variable response in *ilv2.* Constructs encoding the readthrough-promoting cyclic peptides (CPs) were isolated, re-transformed into fresh *S. cerevisiae* IS110-18A cells, and spotted in serial ten-fold dilutions on the relevant selective media. The presence of galactose (gal), but not glucose (gluc), induces CP expression, while the omission of leucine (–Leu) selects for transformed cells. Selection plates with the exclusion of uracil (–Ura) (**A**) or isoleucine (–Ile) (**C**) to select for cells expressing CPs capable of suppressing nonsense mutations in the *ura4* (*ura4-1* TAA-A) and *ilv1* (*ilv1-2* TAA-C) genes, respectively. (**B**) Control plate with no induction of CP expression, but with the omission of uracil to select for nonsense-mutation suppression in the presence of the readthrough-inducing drug G418. Growth enhancement through nonsense-mutation suppression was evaluated by comparison to cells expressing a random cyclic peptide (RCP) or an empty vector (EV). Images were taken after four days (A, C) or three days (B) of incubation and are representative of three independent experiments.

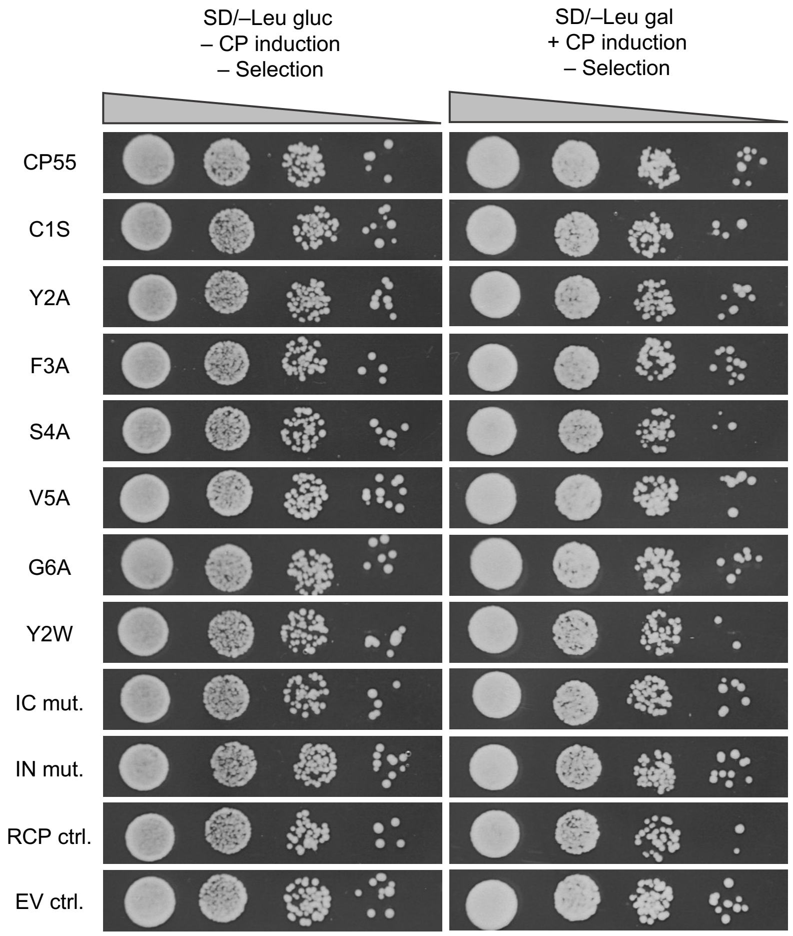


**Figure S5.** CP55 mutants do not affect general *S. cerevisiae* cell growth. Constructs encoding originally selected cyclic peptide (CP) CP55 and derivatives thereof were transformed into *S. cerevisiae* IS110-18A and spotted in serial ten-fold dilutions on control plates. The presence of galactose (gal), but not glucose (gluc) induces CP expression, while omission of leucine (–Leu) selects for transformed cells. Growth was compared to cells expressing a random cyclic peptide (RCP) or an empty vector (EV). Images were taken after two days (left column) or three days (right column) of incubation and are representative of three independent experiments. IC, intein C; IN, intein N.

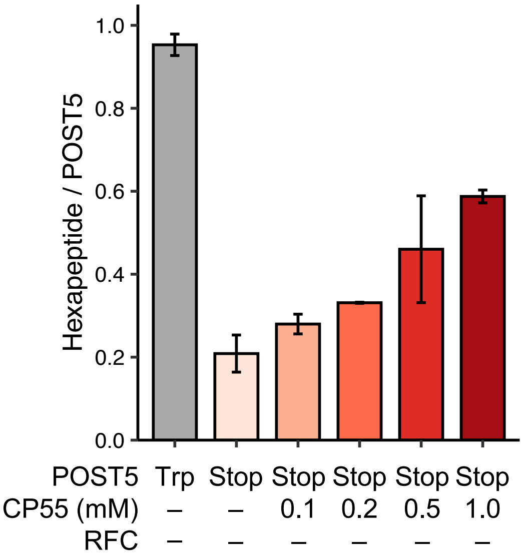


**Figure S6.** CP55 promotes readthrough in co-sedimentation assays in the absence of release factor complex. Readthrough co-sedimentation assay measuring readthrough as the amount of formed hexapeptide relative to POST5 at various concentrations of cyclic peptide (CP) CP55. See Figure 5A for an overview of the experimental principle. A control reaction was included using a POST5 complex with a UGG (Trp) codon in the ribosomal A site, measuring cognate (Trp) amino-acid insertion. The experiment was performed in the absence of the release factor complex (RFC) for comparison with main text Figure 5B in which RFC was included. Shown is the mean ± SD of duplicate reactions.

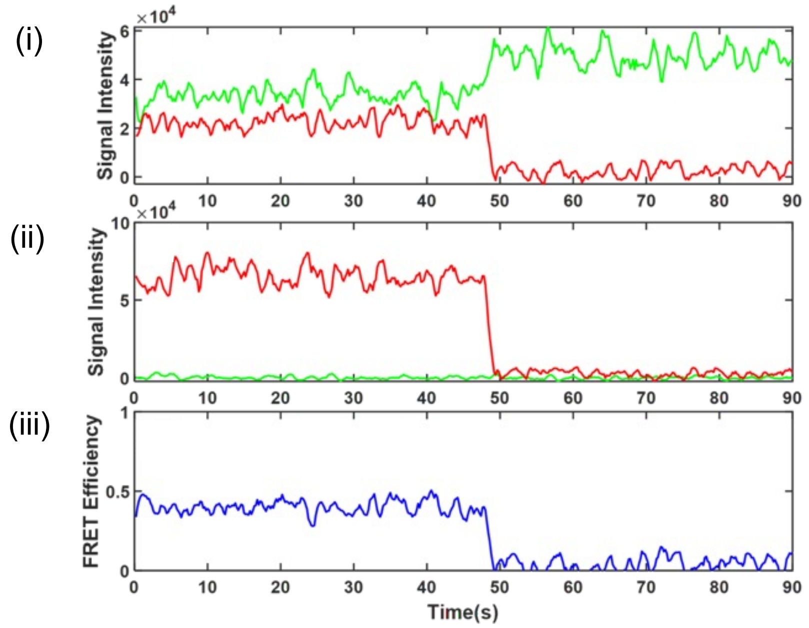


**Figure S7.** Representative recordings from single-molecule FRET microscopy. The traces originate from PRE6 complexes with a Cy5-labeled tRNA in the P site and a Cy3-labeled peptidyl-tRNA in the A site (see Figure 6). (i) Single-molecule intensity recording obtained upon Stop-PRE6 formation in the presence of cyclic peptide (CP) CP55, in which a near-cognate tRNA(Cy3) was bound to the termination codon in the A site. Green and red traces show tRNA(Cy3) donor emission, excited at 532 nm, and tRNA(Cy5) sensitized acceptor emission, respectively, showing anticorrelation and photobleaching of the Cy5 probe at 49 s. (ii) Alternating-laser excitation (ALEX) intensity signal from direct excitation of tRNA(Cy5) at 640 nm. (iii) FRET efficiency between tRNA(Cy3) and tRNA(Cy5) resulting in FRET efficiency of ~0.5 indicative of PRE6 formation.
